# Screening and identification of key biomarkers in diabetic kidney disease and its complications: Evidence from bioinformatics and next generation sequencing data analysis

**DOI:** 10.1101/2023.04.16.537071

**Authors:** Basavaraj Vastrad, Chanabasayya Vastrad

## Abstract

Diabetic patients are prone to diabetic kidney disease (DKD), which may cause cardiovascular damage, hypertension and obesity, and reduce quality of life. As a result, the life quality of patients was seriously reduced. However, the pathogenesis of diabetic kidney disease (DKD) has not been fully elucidated, and current treatments remain inadequate. Therefore, it is essential to explore the molecular mechanism of DKD and its complications. Next Generation Sequancing (GSE217709) dataset was obtained from the Gene Expression Omnibus (GEO) database. Differentially expressed genes (DEGs) were picked out by R software. Then Gene ontology (GO) and REACTOME pathway enrichment analysis were performed by g:Profiler database, protein–protein interaction (PPI) of DEGs was constructed by Human Integrated Protein-Protein Interaction rEference (HIPPIE) database. Module analysis was carried out by Cytoscape plug-in PEWCC. Subsequently, miRNA-hub gene regulatory network and TF-hub gene regulatory network were performed by miRNet database and NetworkAnalyst database. Finally, validation of hub genes was performed by receiver operating characteristic (ROC) curve analysis to predict the diagnostic effectiveness of the hub genes. In total, 958 DEGs, including 479 up regulated and 479 down regulated genes, were identified. The GO and pathway enrichment changes of DEGs were mainly enriched in biological regulation, multicellular organismal process, signaling by GPCR and extracellular matrix organization. Ten hub genes (HSPA8, HSP90AA1, HSPA5, SDCBP, HSP90B1, VCAM1, MYH9, FLNA, MDFI and PML) associated with DKD and its complications were identified. Bioinformatics analysis is a useful tool to explore the molecular mechanism and pathogenesis of DKD and its complications. The identified hub genes may participate in the onset and development of DKD and its complications and serve as therapeutic targets.

## Introduction

Diabetic kidney disease (DKD) is a microvascular diseases in diabetes mellitus and it affects 40% of people with type 2 diabetes mellitus and 30% of those with type 1 diabetes mellitus worldwide [1–2]. DKD represent the leading cause of chronic kidney disease and end-stage renal disease [3]. Like chronic kidney disease, there is a significant risk of morbidity and mortality, as well as a decreased quality of life and an increase in healthcare costs associated with DKD [4]. DKD was characterized by renal hemodynamics changes, oxidative stress, inflammation, hypoxia, overactive renin-angiotensin-aldosterone system and renal fibrosis [5]. The number of cases of DKD rising globally and it has become a key health concern [6]. These patients have higher risks of cardiovascular damage [7], hypertension [8] and obesity [9]. DKD displays a high heterogeneity, which is strongly linked with morphology characteristics and molecular changes [10]. Hence, developing novel diagnosis biomarkers reflecting the damage of kidney is appealing for the timely diagnoses and therapies of DKD.

Despite a number of genes and signaling pathways in the development and progression of DKD have been widely investigated, the mechanisms underlying DKD are still being unraveled. With the advancement of next generation sequencing (NGS) technology, integrated bioinformatics provides an effective tool for discovering valuable new biomarker targets. Recently, many specific genes have been detected to participate in the advancement of DKD. LRG1 [11], STAT1 [12], phosphofructokinase platelet type (PFKP) [13], ADIPOQ (adiponectin) [14] and COL4A3 [15] are associated with DKD. Signaling pathways include Wnt/β-catenin signaling pathway [16], AMPK/SIRT1-FoxO1 signaling pathway [17], PI3K/Akt/mTOR signaling pathway [18], JAK/STAT signaling pathway [19] and PI3K/Akt/NF-κB signaling pathway [20] and ROS/NLRP3 signaling pathway [21] were found to be substantially related to DKD. These finding suggested the essential roles of some function genes in DKD advancement. However, the diagnostic value of many genes has not been studied in DKD.

NGS technology which was used recently can quickly detect differentially expressed genes (DEGs) and was proved to be a reliable technique that could make data be produced and stored in public databases [22]. Therefore, a large number of valuable clues could be explored for new investigation on the base of these NGS data. Furthermore, many bioinformatics studies on DKD have been produced in recent years [23], which proved that the integrated bioinformatics methods could help us to further study and better exploring the underlying molecular mechanisms.

In the current investigation, we selected NGS dataset GSE217709 from the Gene Expression Omnibus (GEO) database (https://www.ncbi.nlm.nih.gov/geo/) [24], and used the DESeq2 package in R software [25] to screen DEGs. Subsequently, we analyzed the Gene Ontology (GO) functions and REACTOME pathways associated with the resulting DEGs. Moreover, a protein-protein interaction (PPI) network, modules, miRNA-hub gene regulatory network and TF-hub gene regulatory network were established, and hub genes, miRNAs and TFs were selected. Then, we analyzed their diagnostic value of hub genes in DKD based on receiver operating characteristic (ROC) curve. Our results might provide potential and novel biomarker candidates for clinical diagnosis and therapy of DKD.

## Materials and Methods

### Next generation sequencing data source

The NGS data of GSE217709, downloaded from GEO database were used to screen DEGs in DKD. 38 specimens (29 DKD samples and 9 normal control samples) were included in the data series of GSE217709. And the gene expression was detected by GPL20301 Illumina HiSeq 4000 (Homo sapiens).

### Identification of DEGs

The analysis tool DESeq2 was using to identify the DEGs. DEGs were selected with threshold of log2FC > 0.58, log2FC < -1.044 and adj p value < 0.05. DEGs with log2FC > 0.58 were considered as up regulated genes, and log2FC < -1.044 as down regulated genes. Volcano plots and heatmap for DEGs were created via ggplot2 and gplot packages of R software.

### GO and pathway enrichment analyses of DEGs

To get a better understanding of the function of DEGs, g:Profiler (http://biit.cs.ut.ee/gprofiler/) [26] was used to conduct REACTOME (https://reactome.org/) [27] pathway enrichment analyses of common DEGs. GO enrichment analysis (GO, http://www.geneontology.org) [28] consisting of biological process (BP), cellular component (CC), and molecular function (MF) were also performed. The statistically significant enriched BP, CC, and MF terms and all enriched REACTOME pathways were recorded. g:Profiler was used to explore GO enrichment and REACTOME pathway enrichment analysis with a cut-off criterion of adjusted P<0.05.

### Construction of the PPI network and module analysis

In order to obtain directly or indirectly interacting proteins related to DEGs, the Human Integrated Protein-Protein Interaction rEference (HIPPIE) (http://cbdm-01.zdv.uni-mainz.de/~mschaefer/hippie/) PPI database was used [29]. Then, PPI network visualization is constructed by using Cytoscape software (http://www.cytoscape.org/) [30]. The Network Analyzer plugin of Cytoscape (v 3.9.1) was used to score each node gene by 4 selected algorithms, including degree [31], betweenness [32], stress [33] and closeness [34]. Subsequently, the Cytoscape plug-in the PEWCC [35] was used to identify the significant modules from PPI network.

### Construction of the miRNA-hub gene regulatory network

The miRNet database (https://www.mirnet.ca/) [36] is an open-source platform mainly focusing on miRNA-hub gene interactions. miRNet utilizes 14 established miRNA-hub gene prediction databases, including TarBase, miRTarBase, miRecords, miRanda (S mansoni only), miR2Disease, HMDD, PhenomiR, SM2miR, PharmacomiR, EpimiR, starBase, TransmiR, ADmiRE, and TAM 2.0. Subsequently, the miRNA-hub gene regulatory network of the hub genes and their targeted miRNAs was visualized by Cytoscape software [30].

### Construction of the TF-hub gene regulatory network

The NetworkAnalyst database (https://www.networkanalyst.ca/) [37] is an open-source platform mainly focusing on TF-hub gene interactions. NetworkAnalyst utilizes one established TF-hub gene prediction CHEA database. Subsequently, the TF-hub gene regulatory network of the hub genes and their targeted TFs was visualized by Cytoscape software [30].

### Receiver operating characteristic curve (ROC) analysis

To evaluate the role of candidate genes in the diagnosis of DKD, receiver operating characteristic (ROC) curve analysis was conducted in RStudio with pROC package [38]. The calculation of the area under the curve (AUC) was conducted. Thus, we investigated the feasibility of the hub genes for prediction using the AUC value. The genes with area under curve (AUC) >0.9 were considered as hub genes of DKD.

## Results

### Identification of DEGs

In our study, 958 DEGs were identified between DKD samples and normal control samples. Among them, 479 were up regulated genes (log2 FC>0.58) and 479 were down regulated genes (log2 FC< -1.044) (Table 1). The volcano plot and heatmap of gene expression are shown in Fig. 1 and Fig. 2.

**Table 1.**
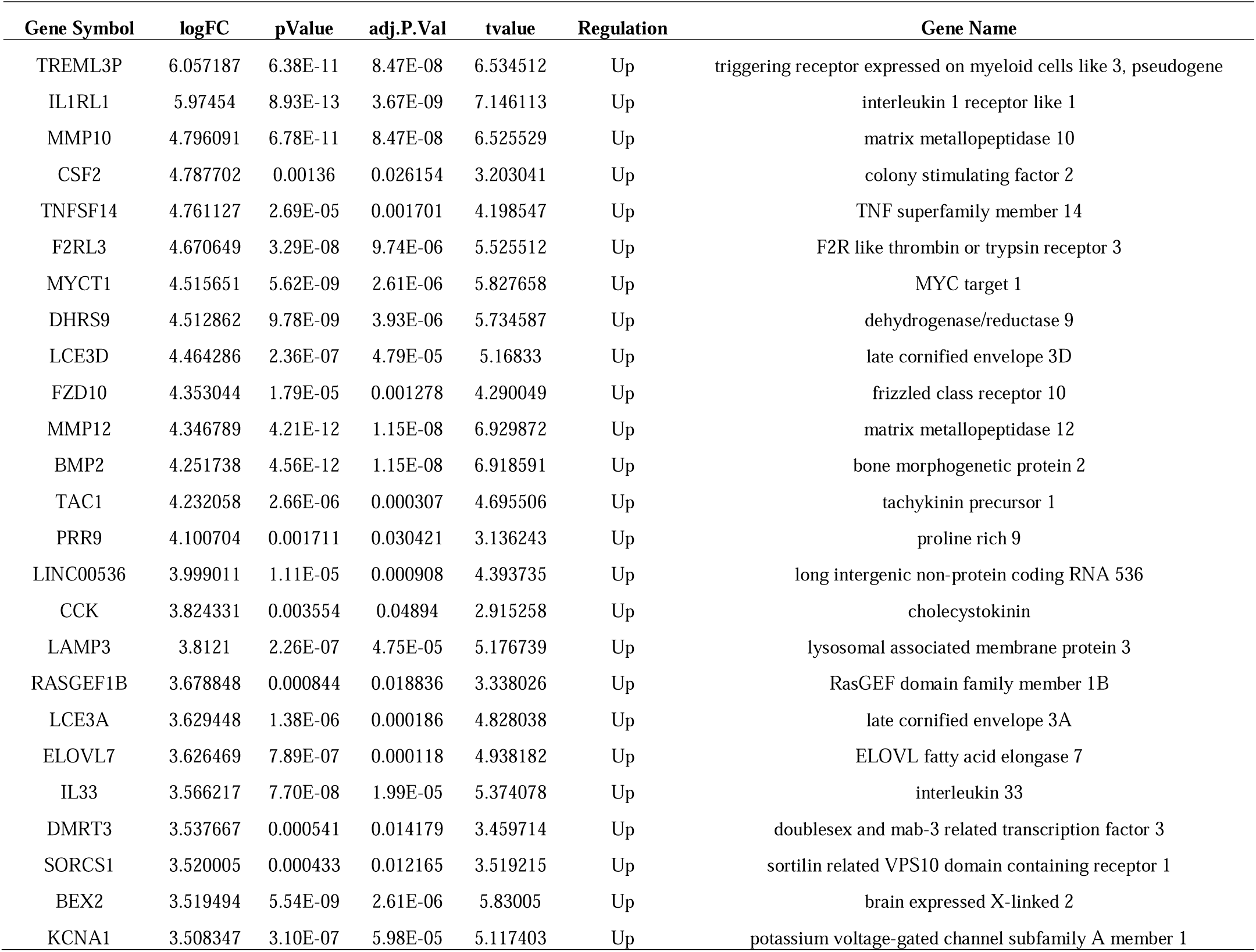
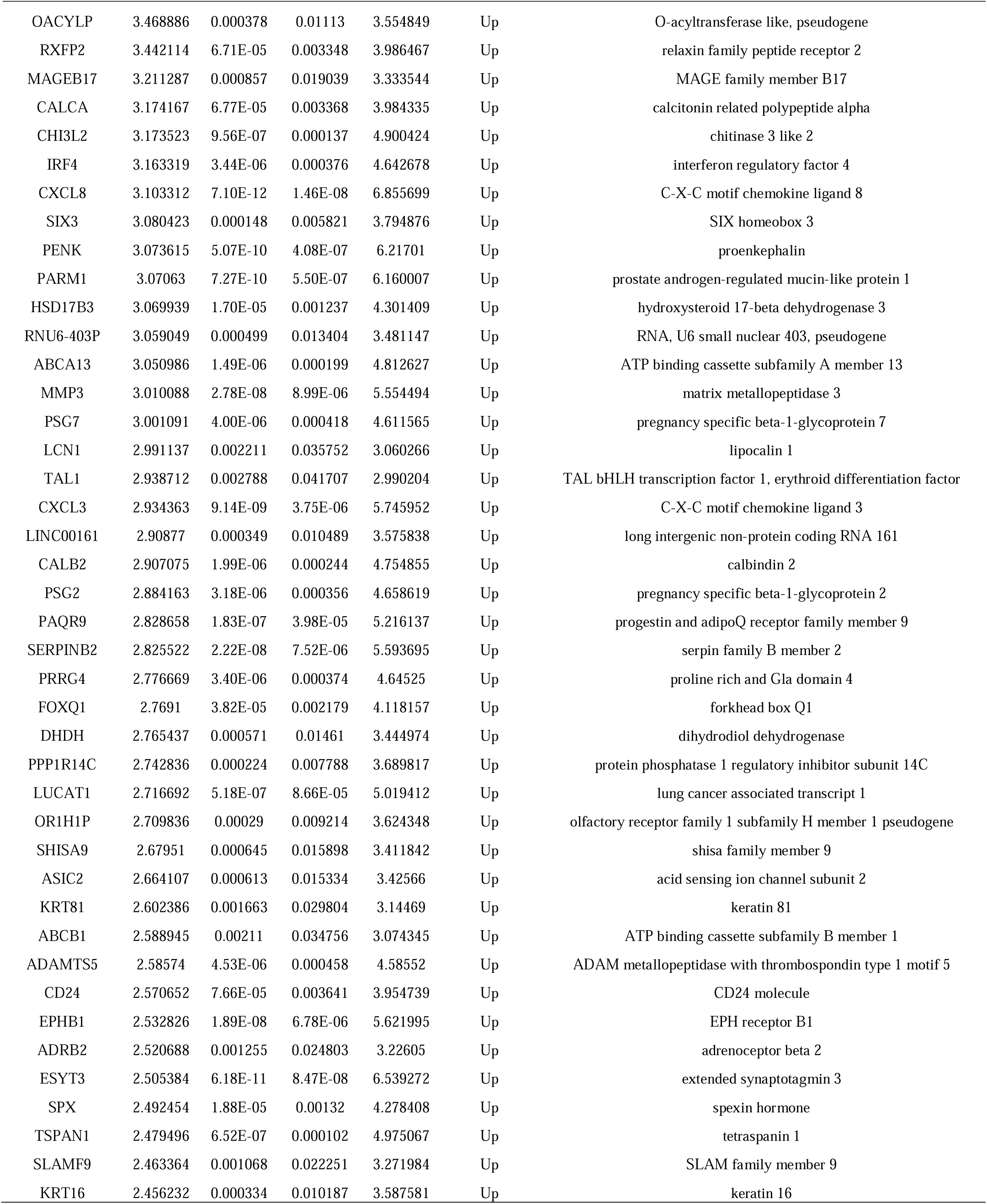
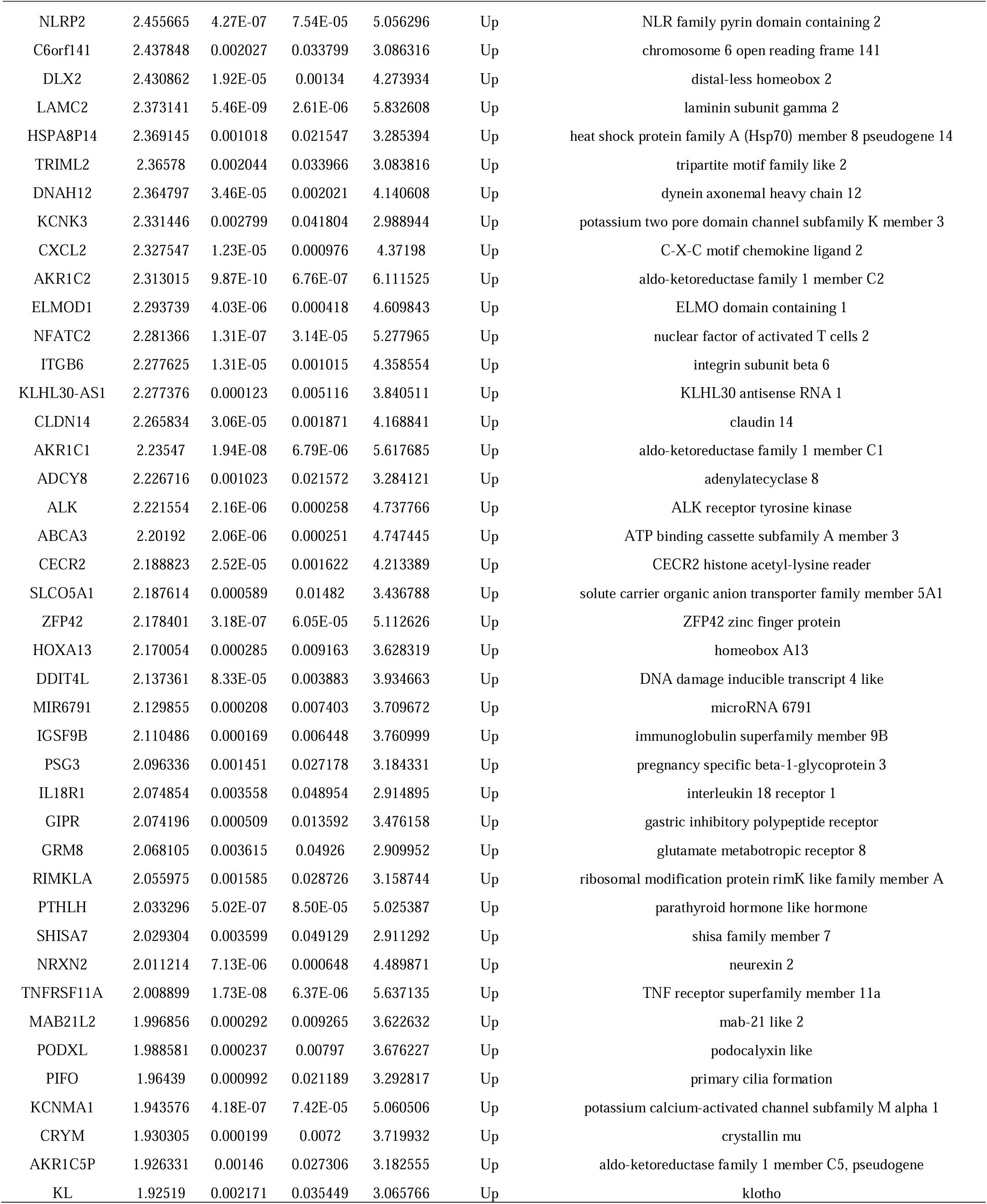
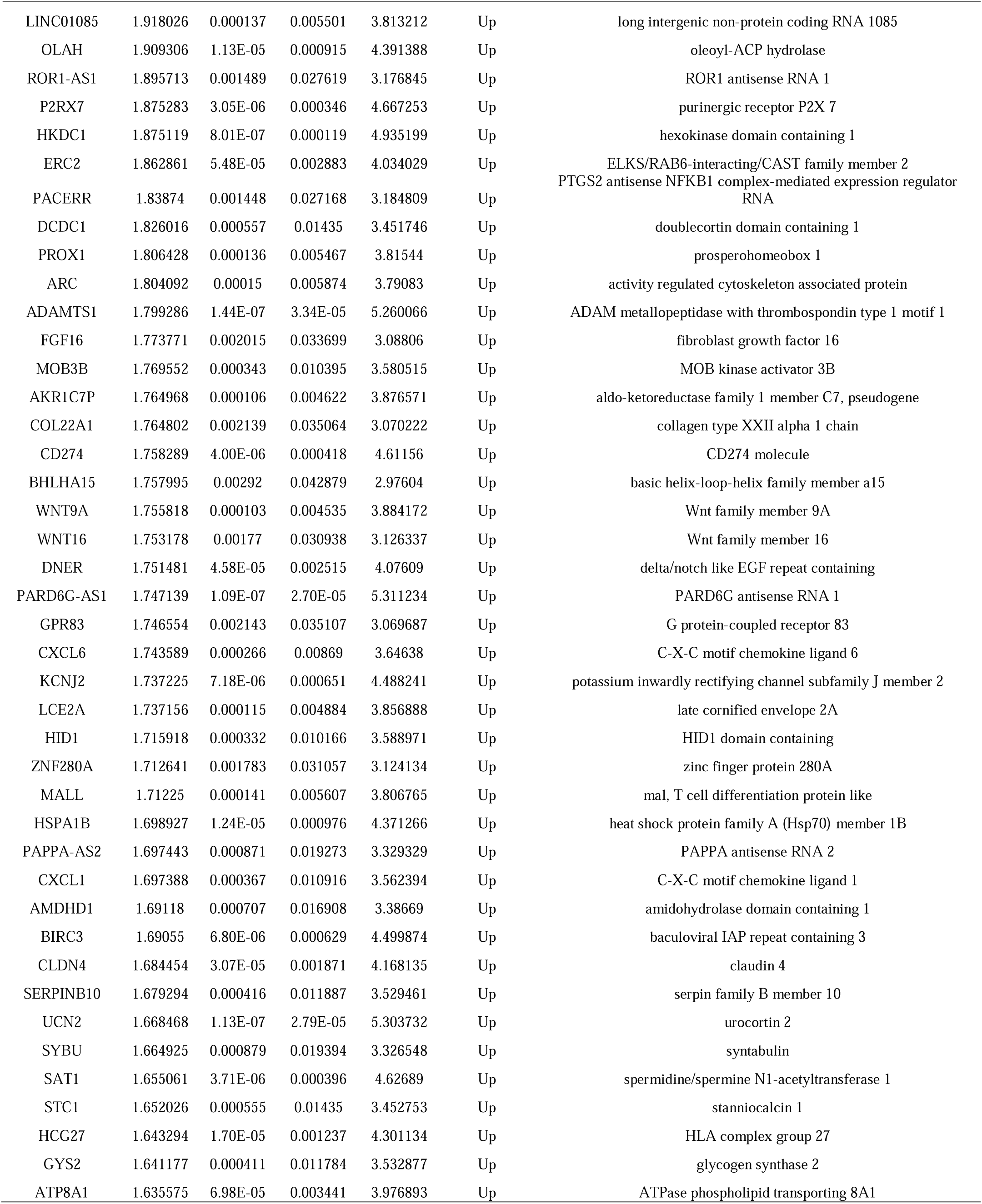
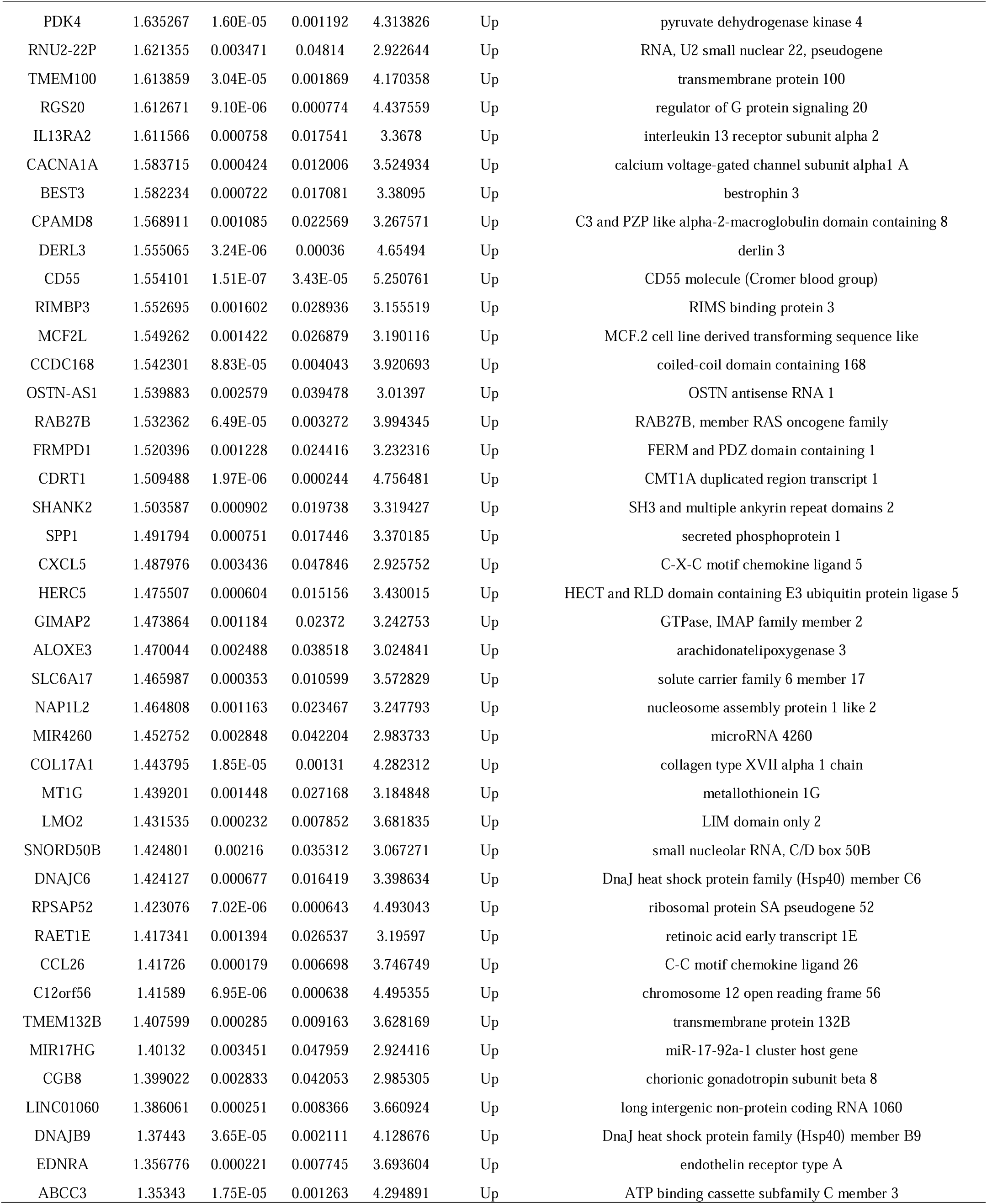
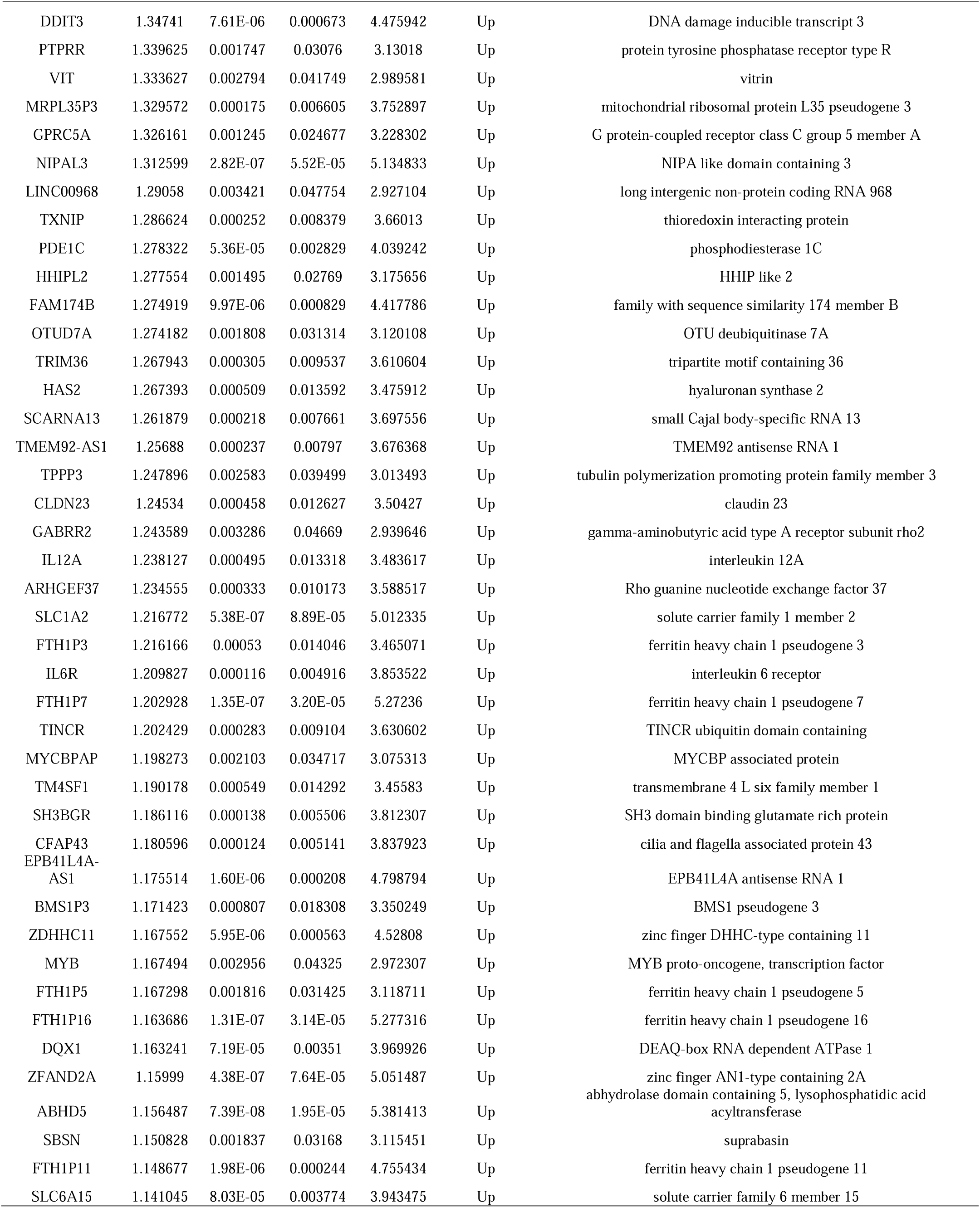
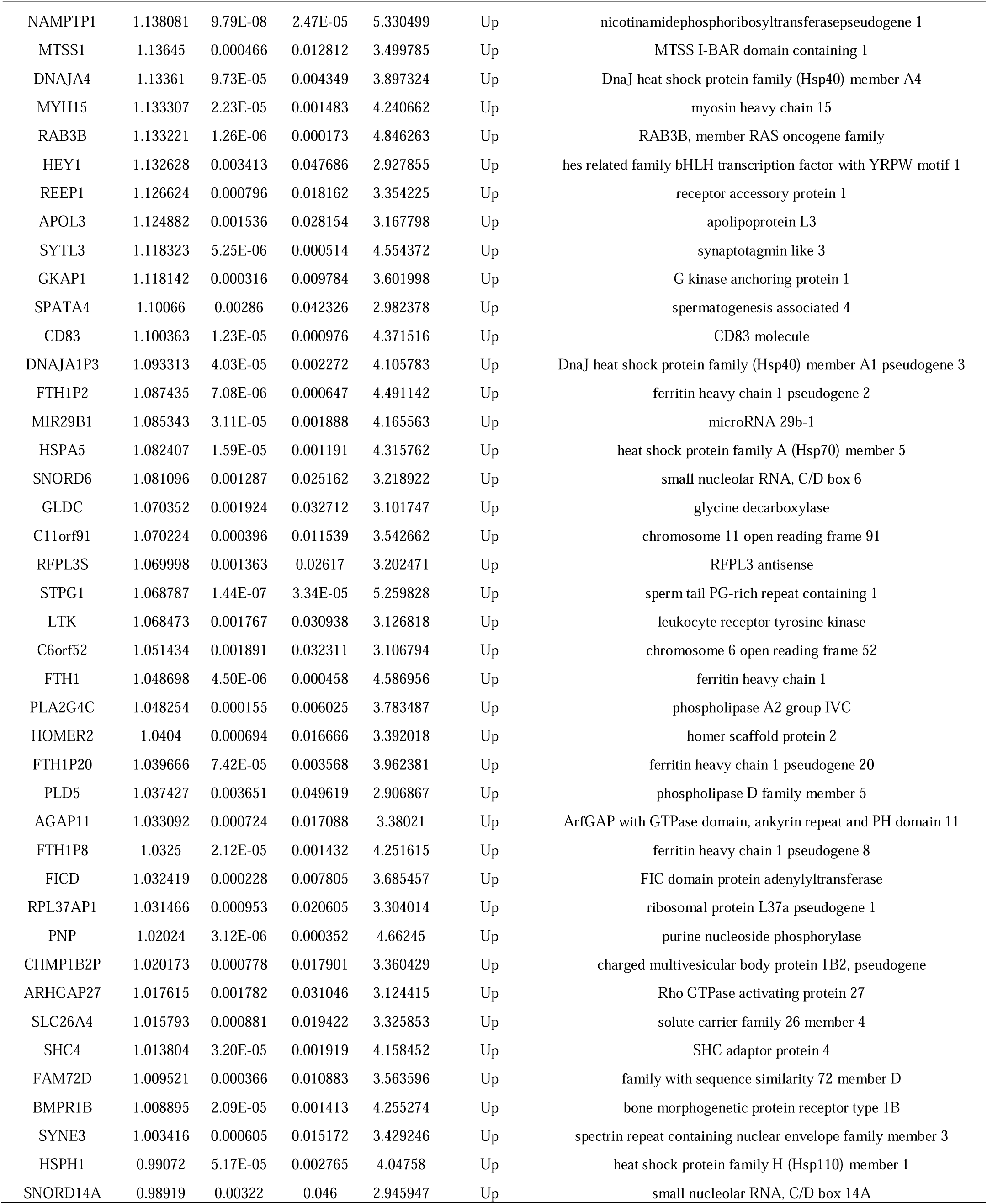
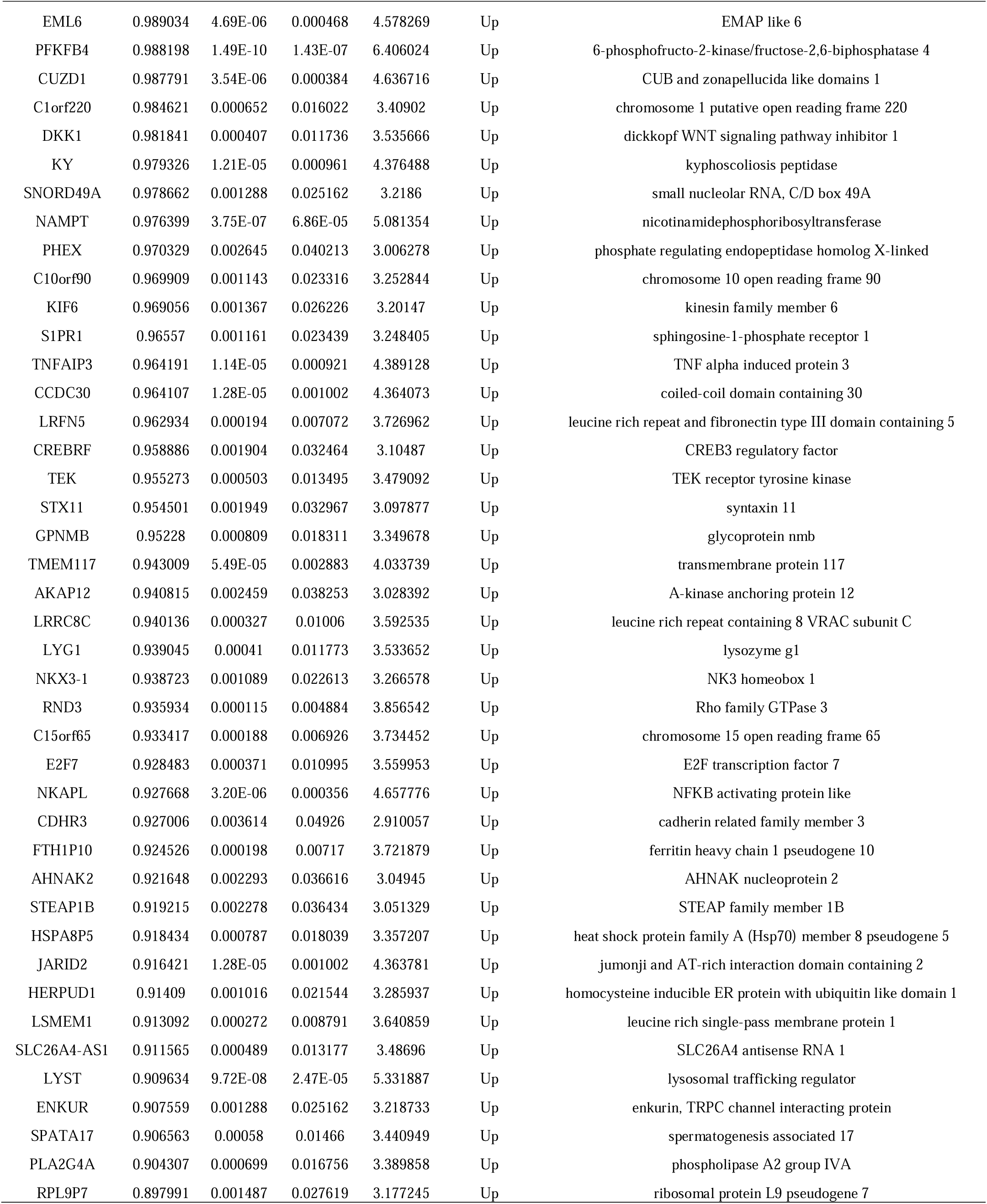
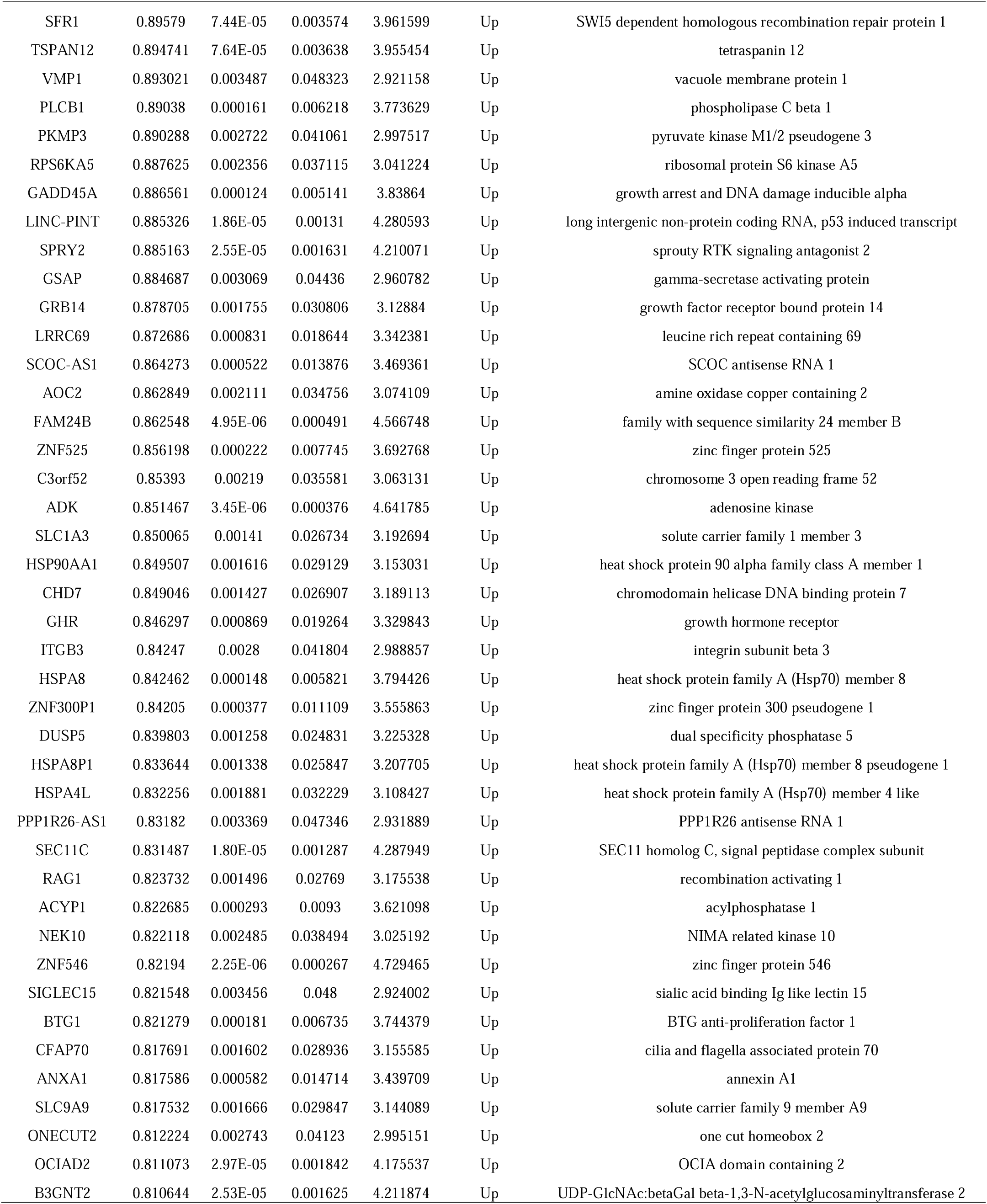
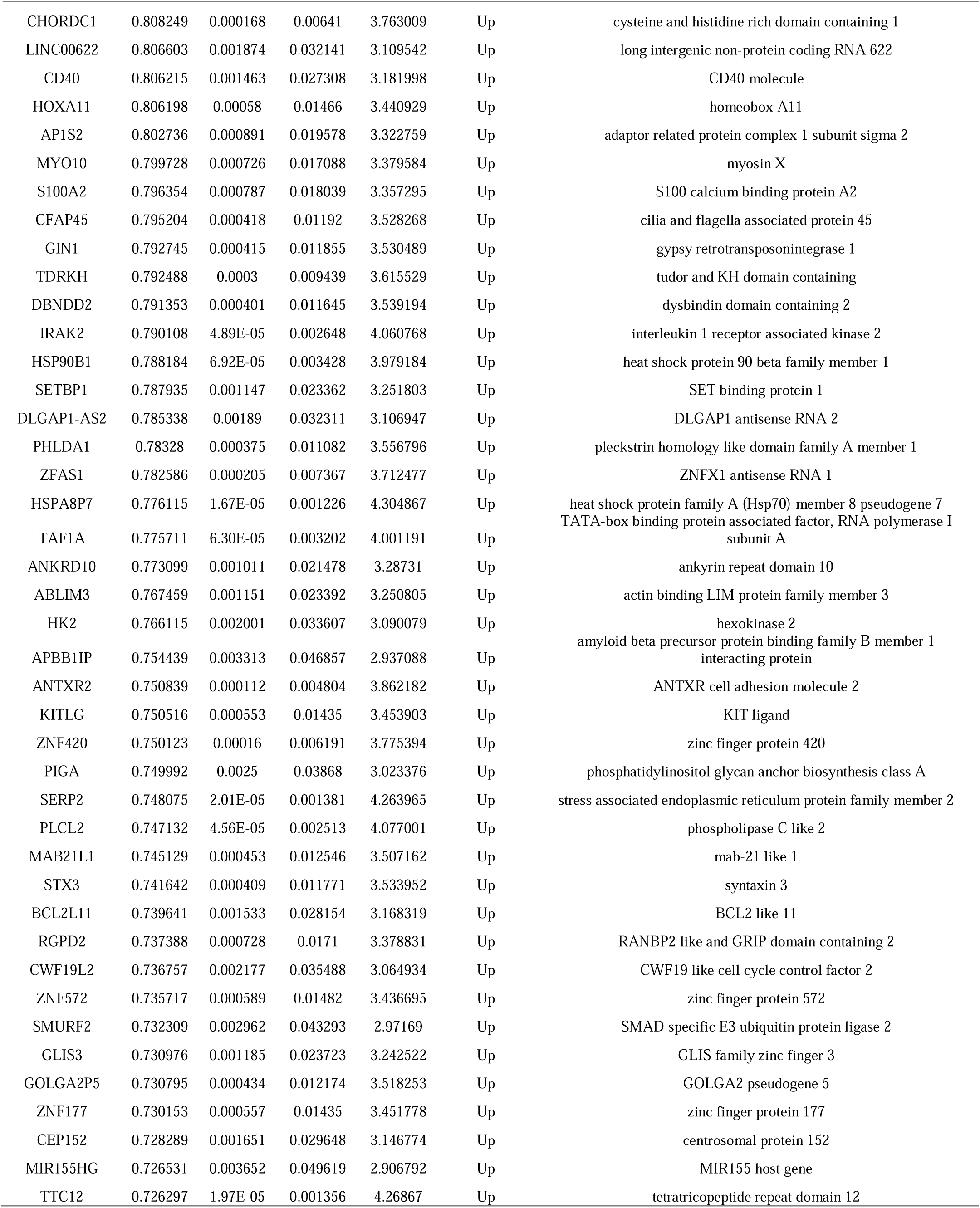
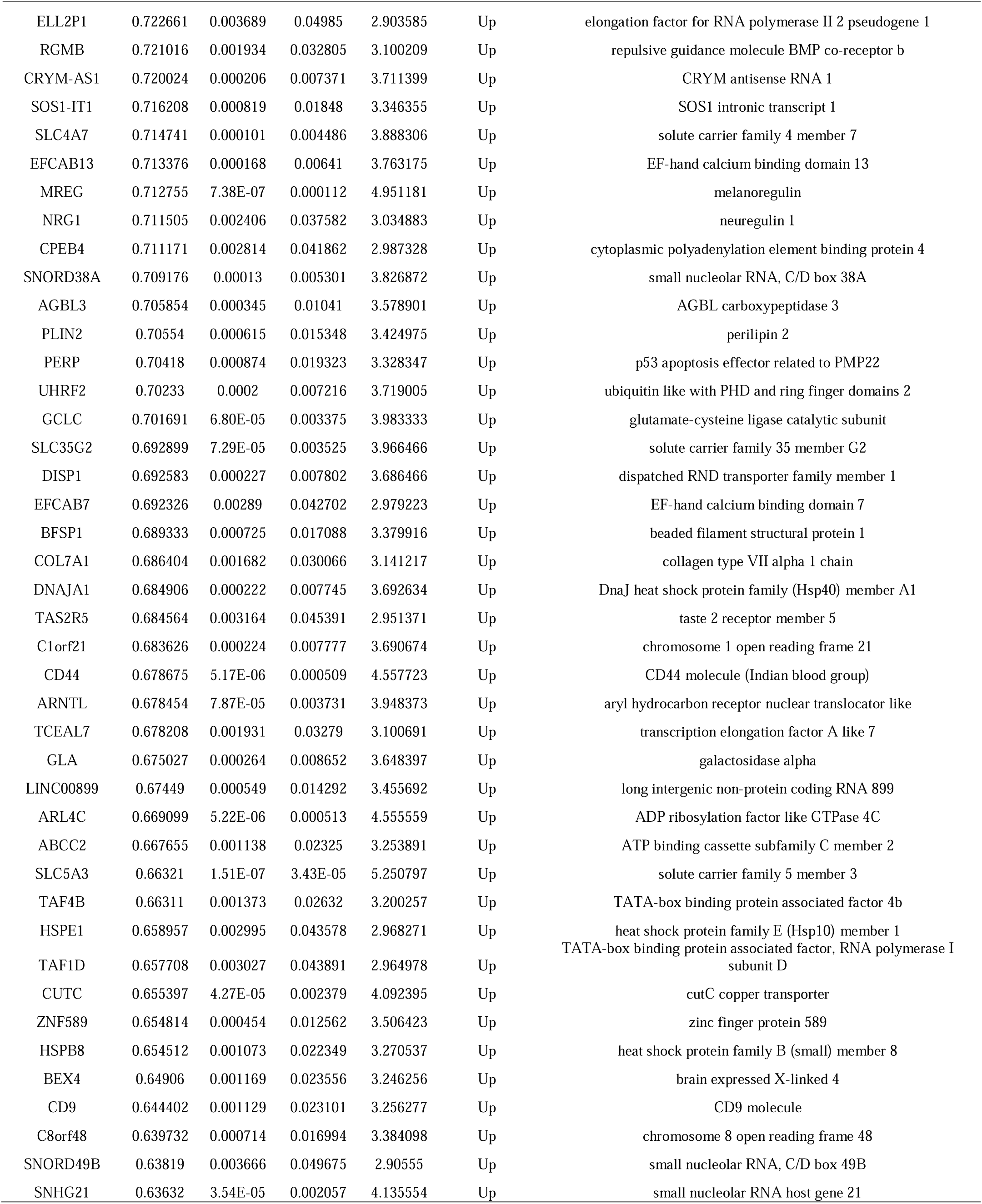
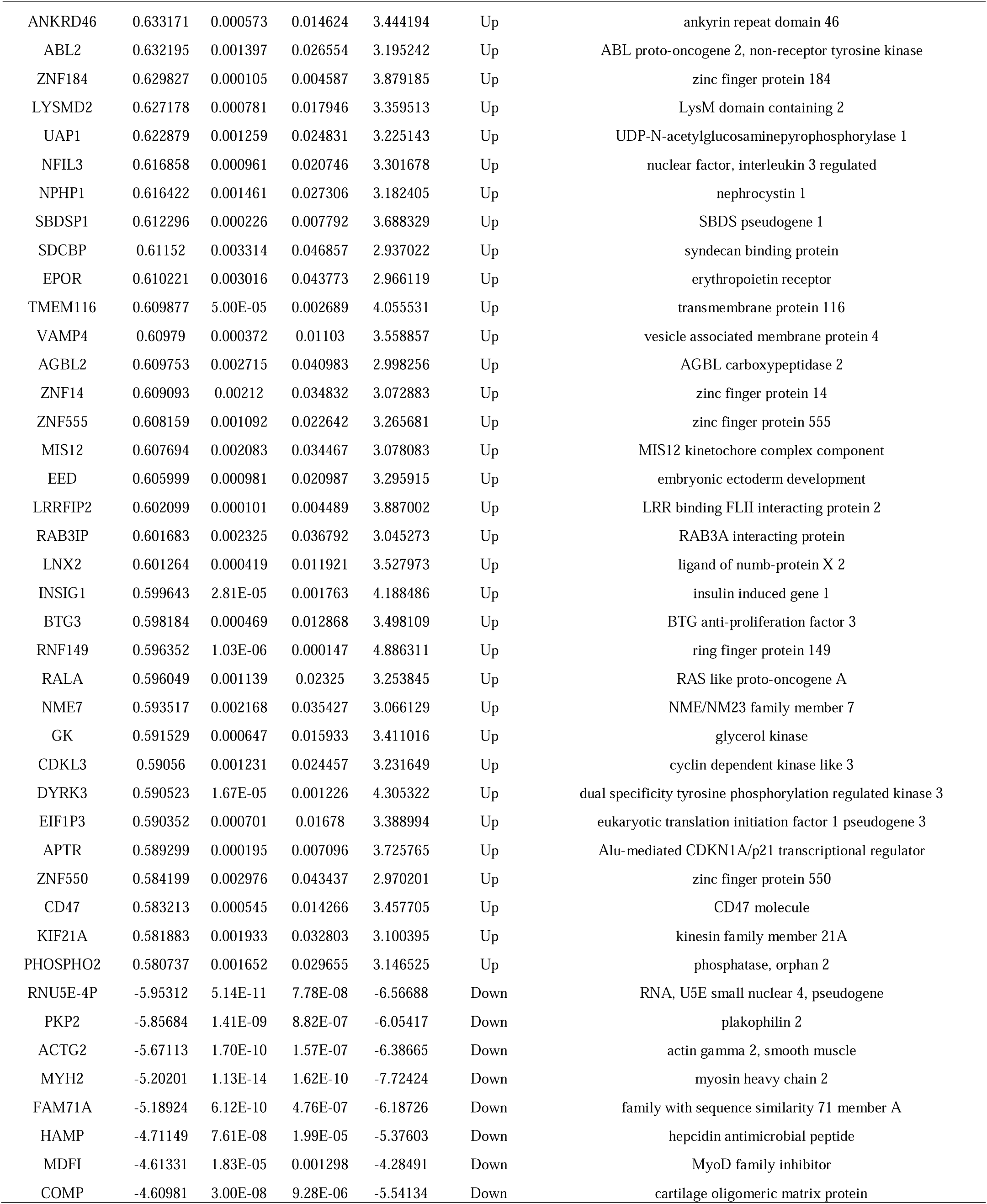
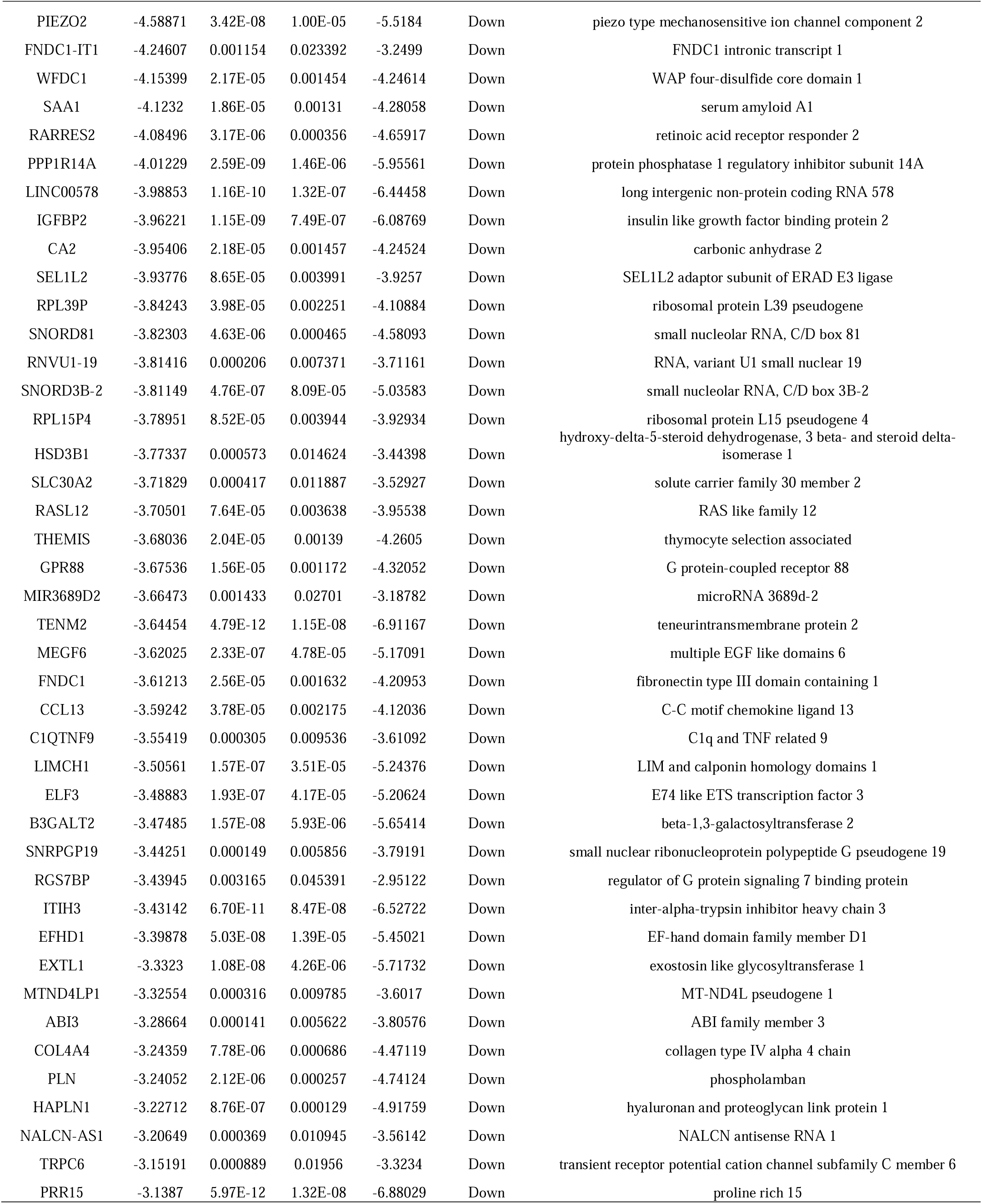
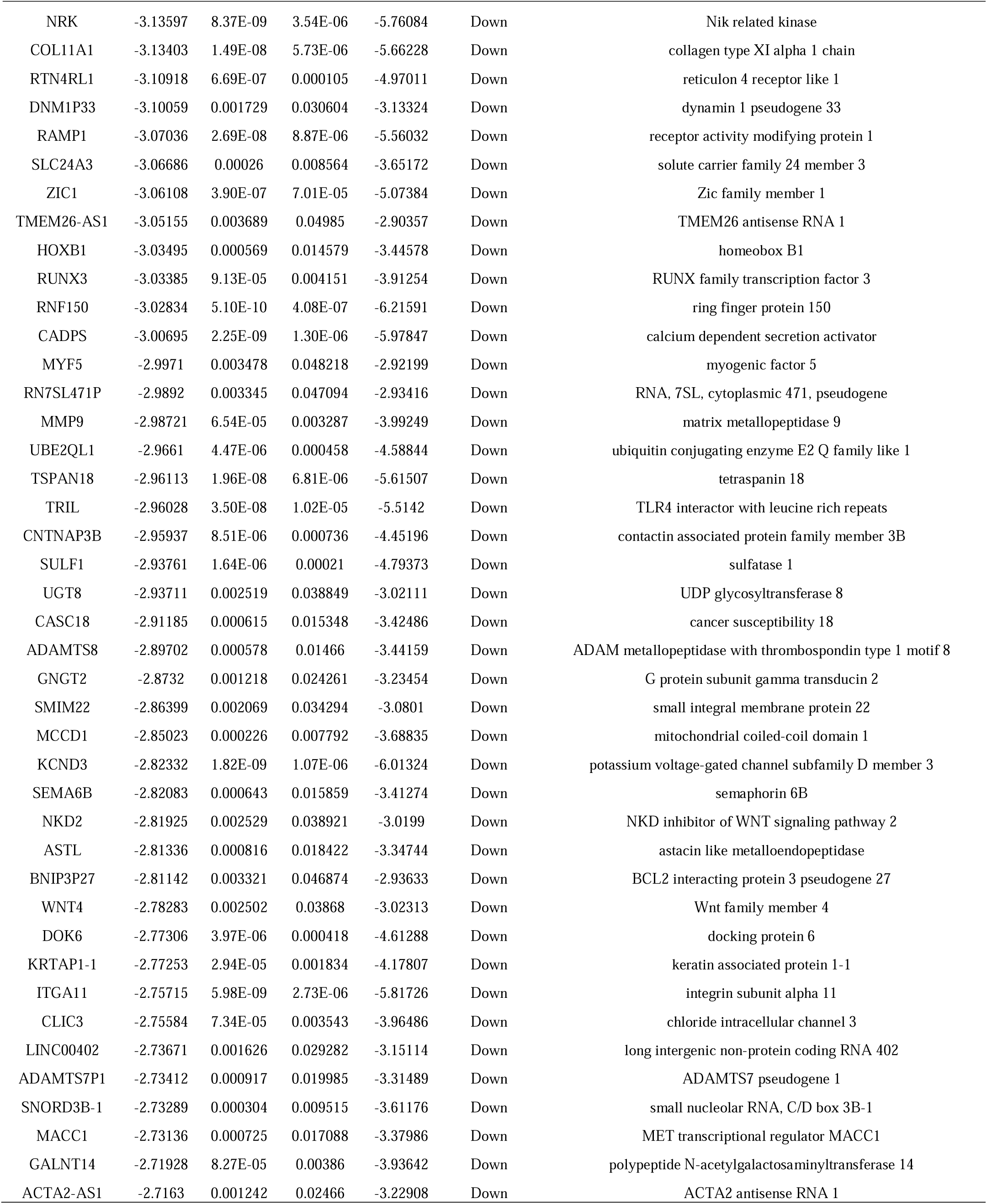
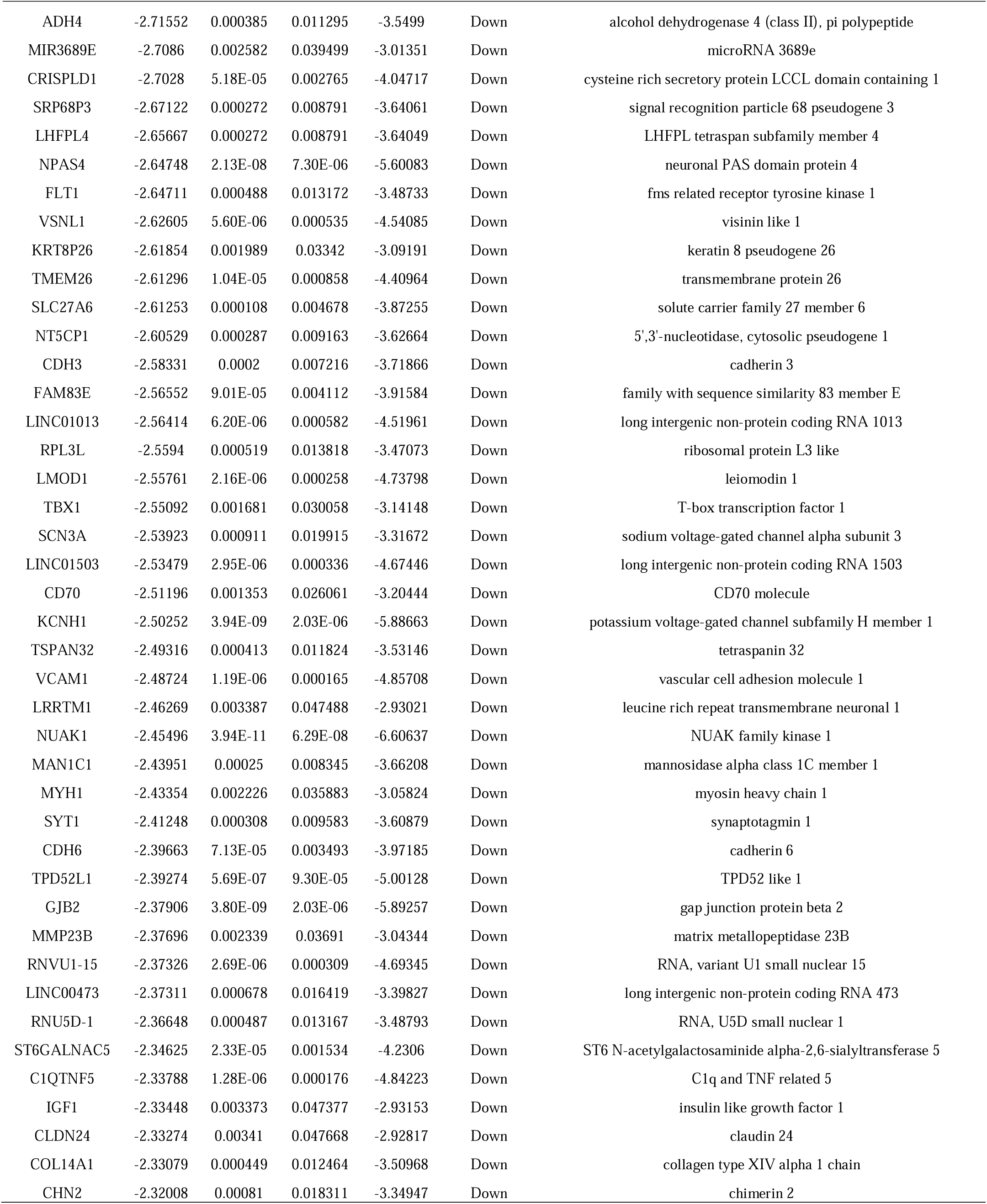
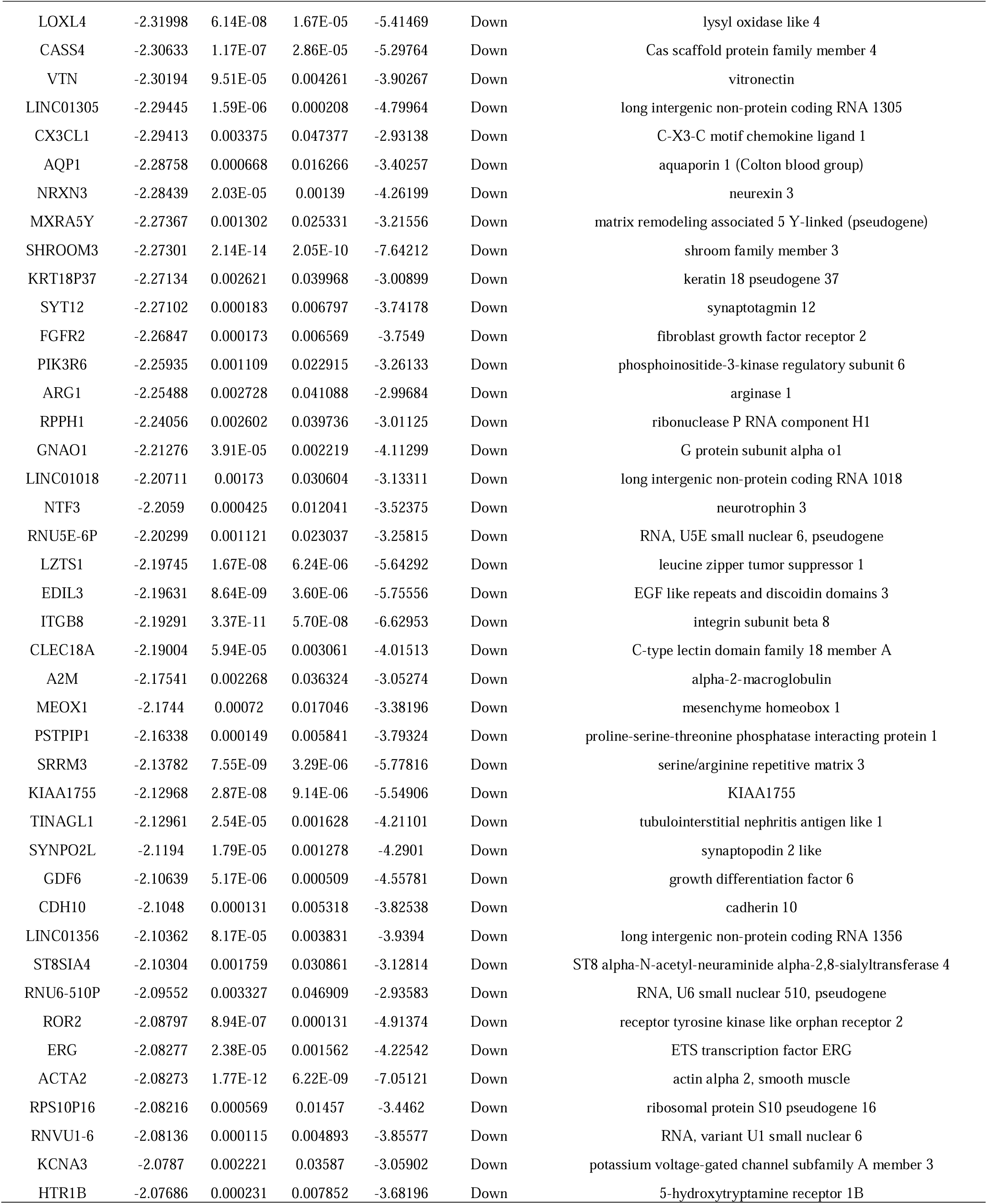
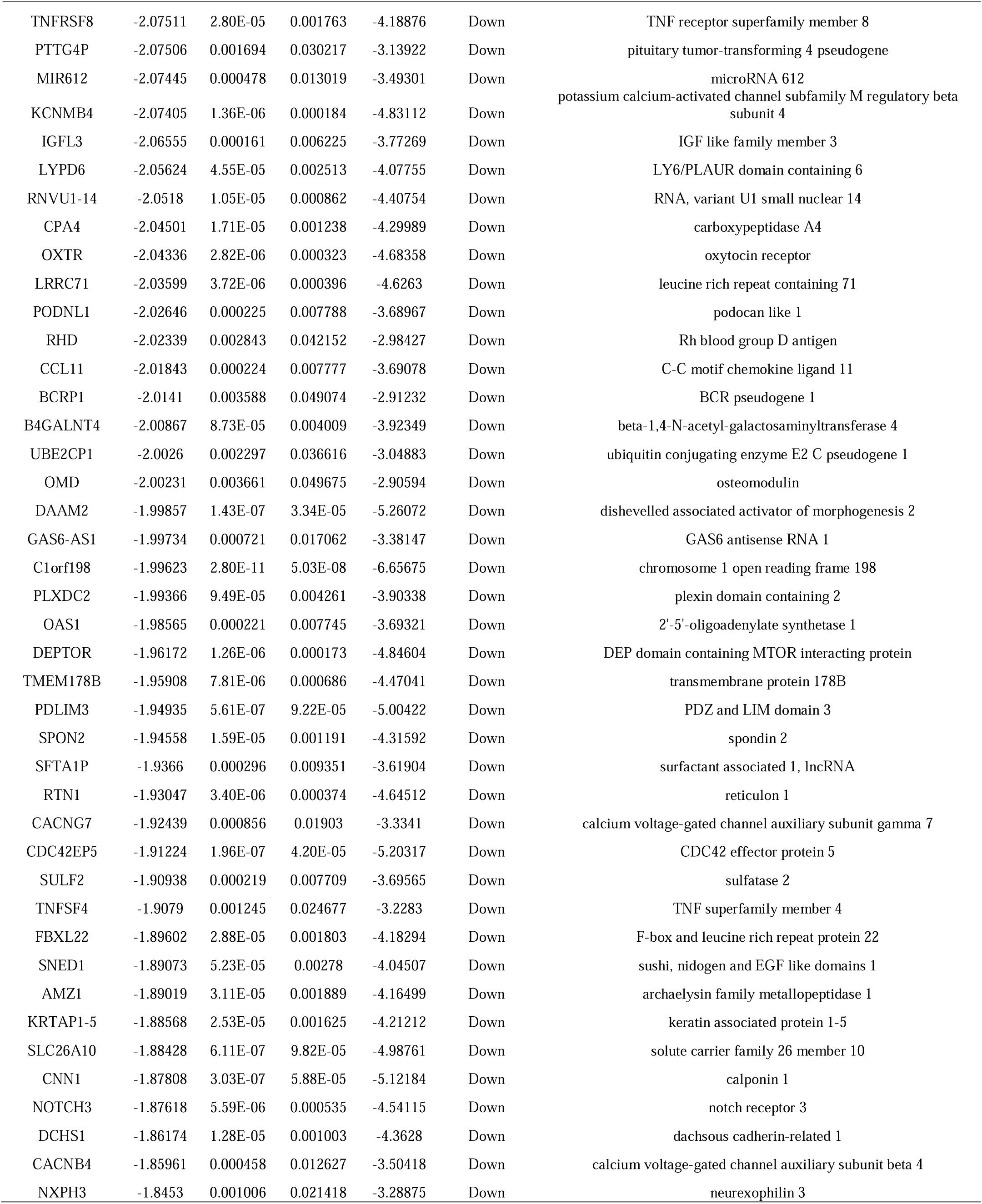
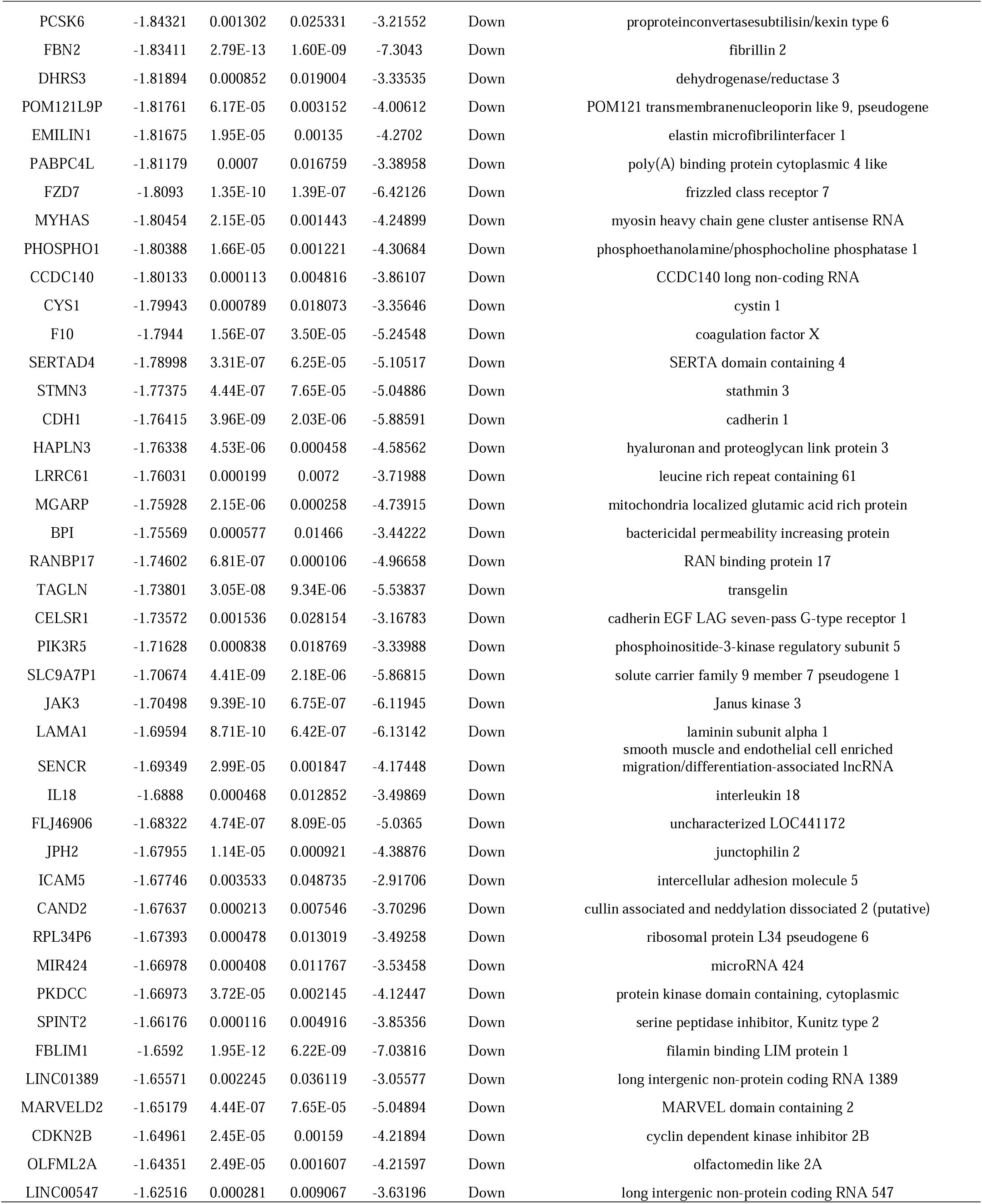
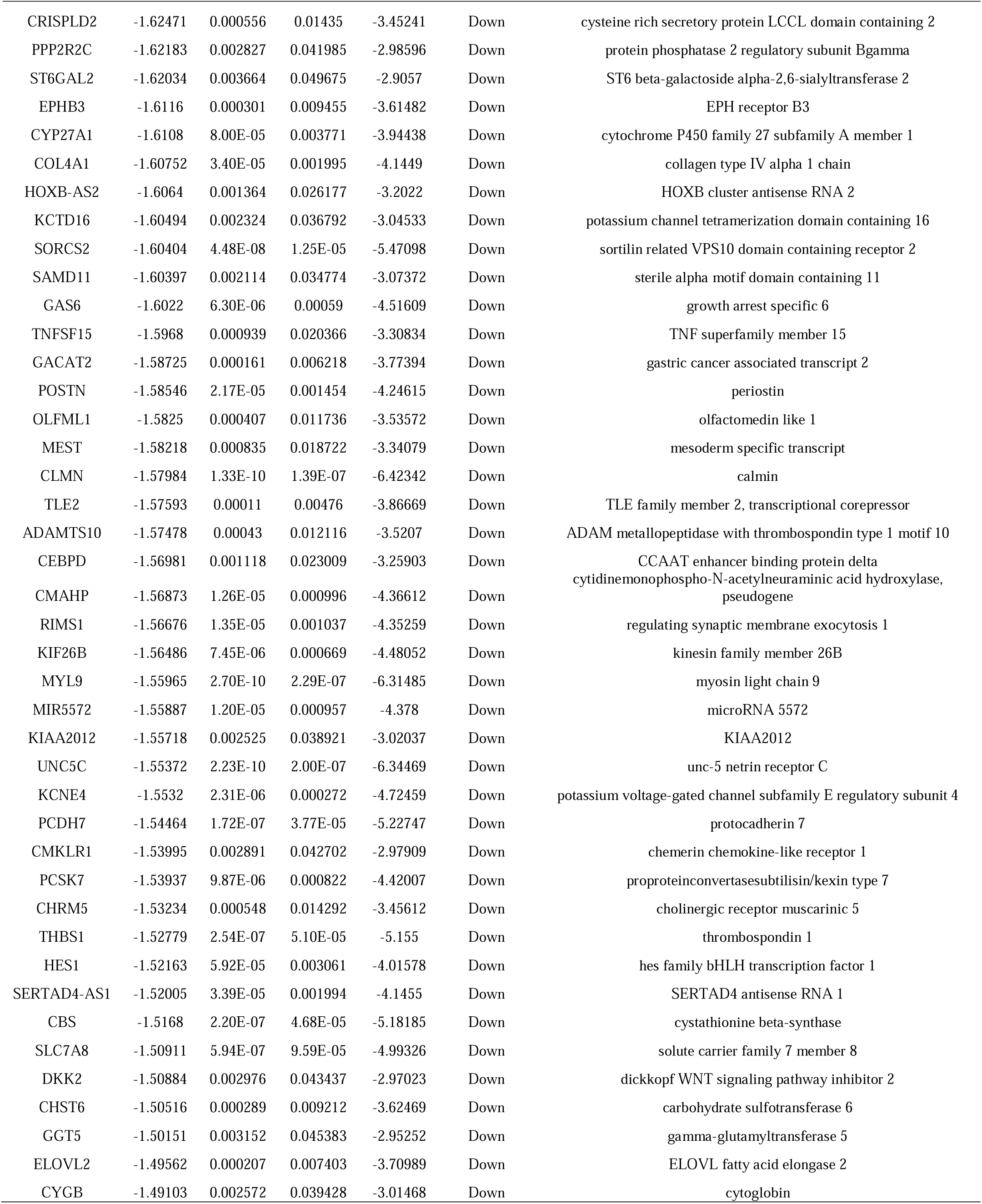
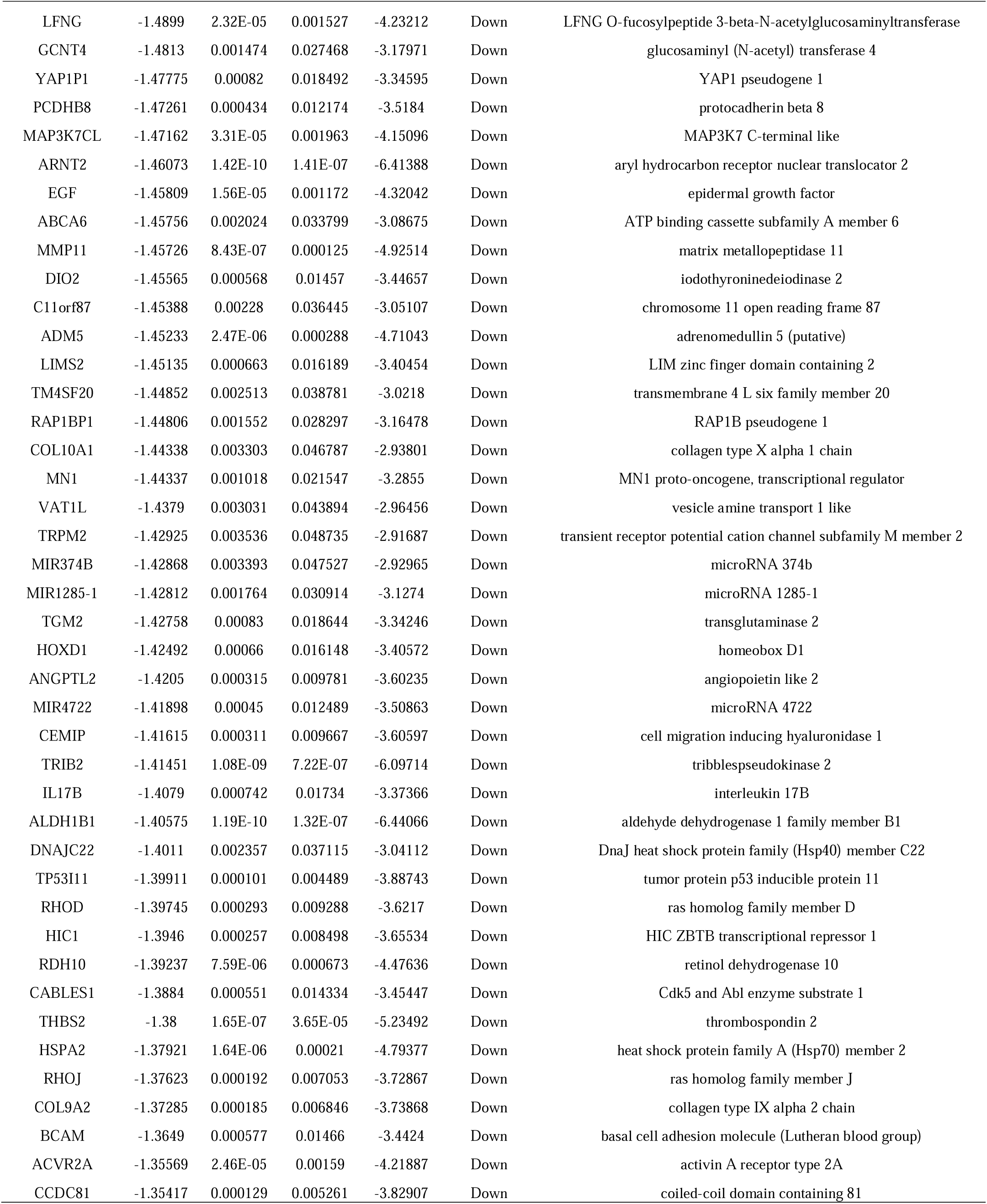
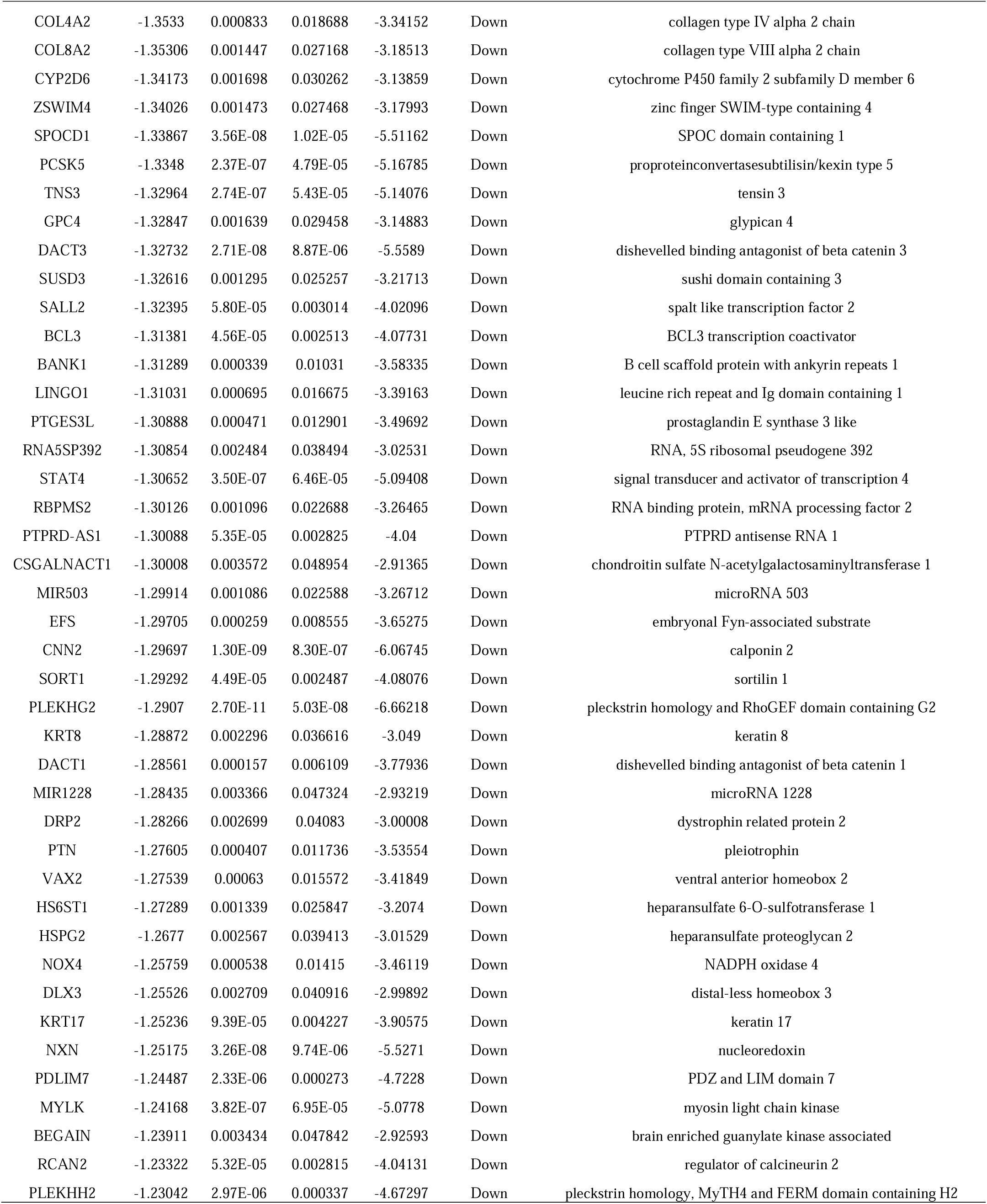
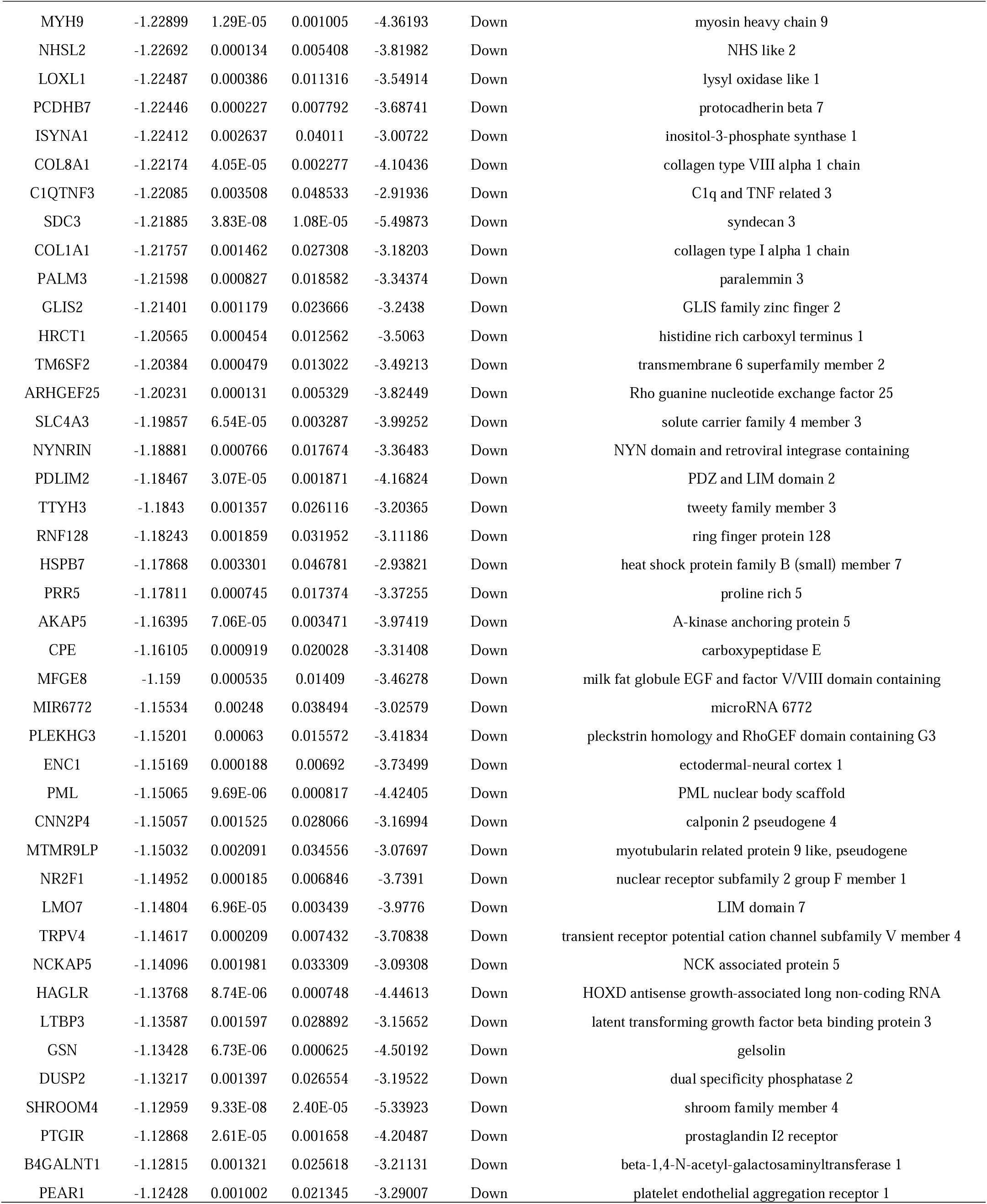
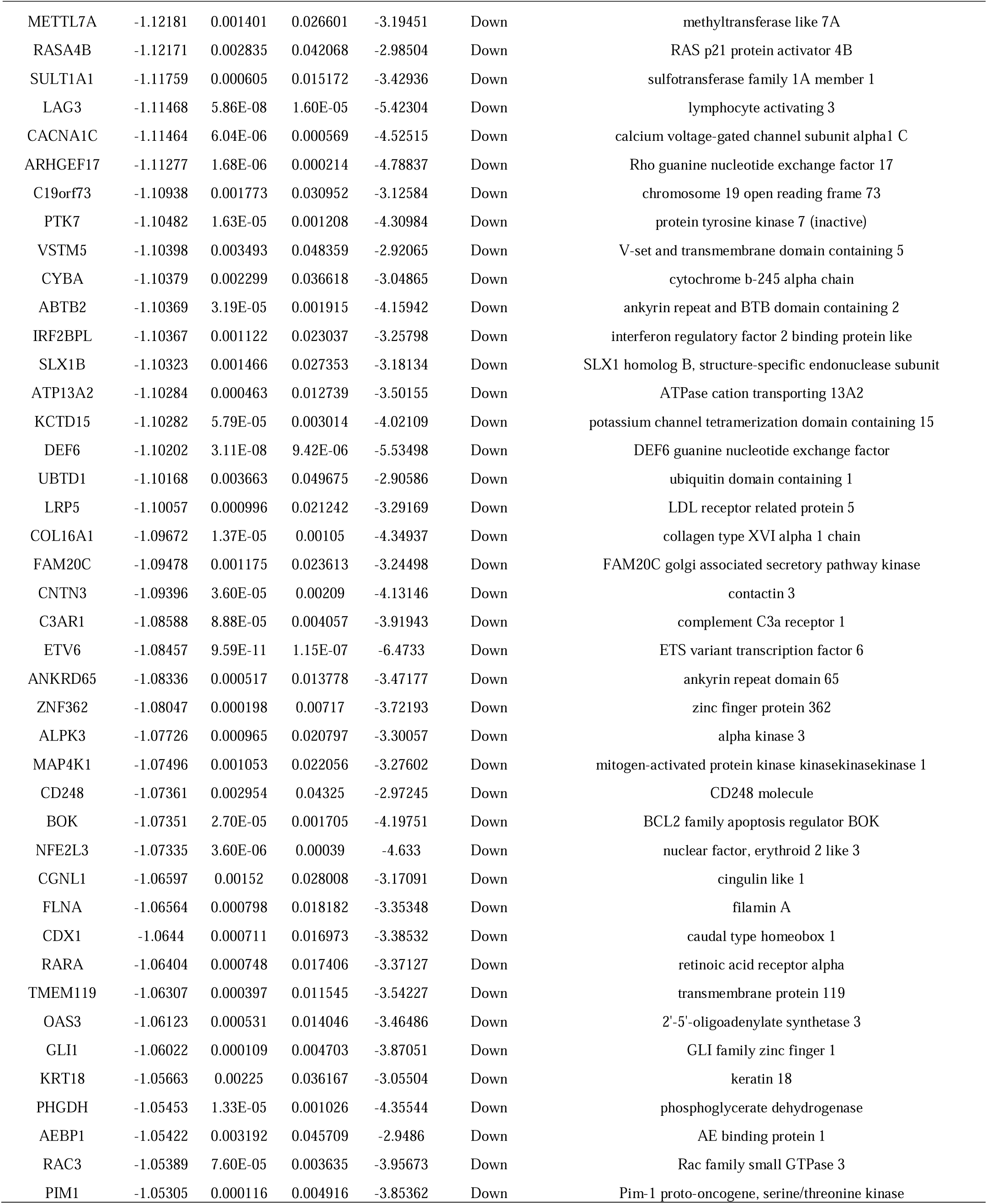
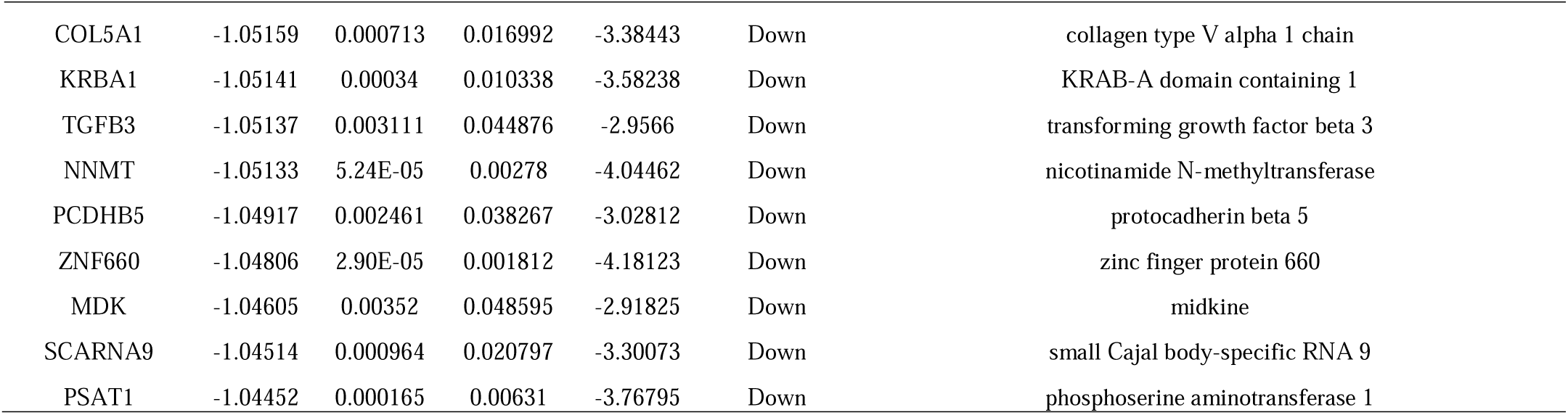
The statistical metrics for key differentially expressed genes (DEGs)

**Fig. 1.**
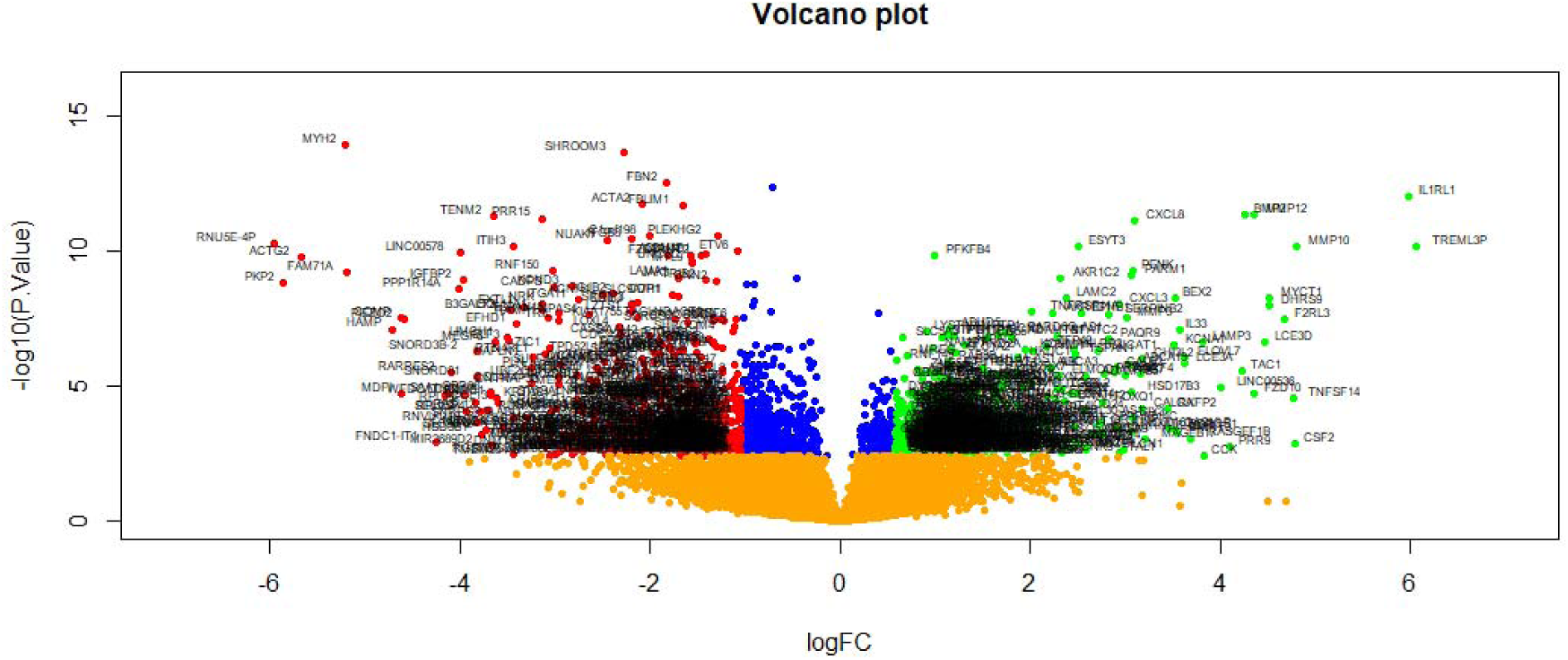
Volcano plot of differentially expressed genes. Genes with a significant change of more than two-fold were selected. Green dot represented up regulated significant genes and red dot represented down regulated significant genes.

**Fig. 2.**
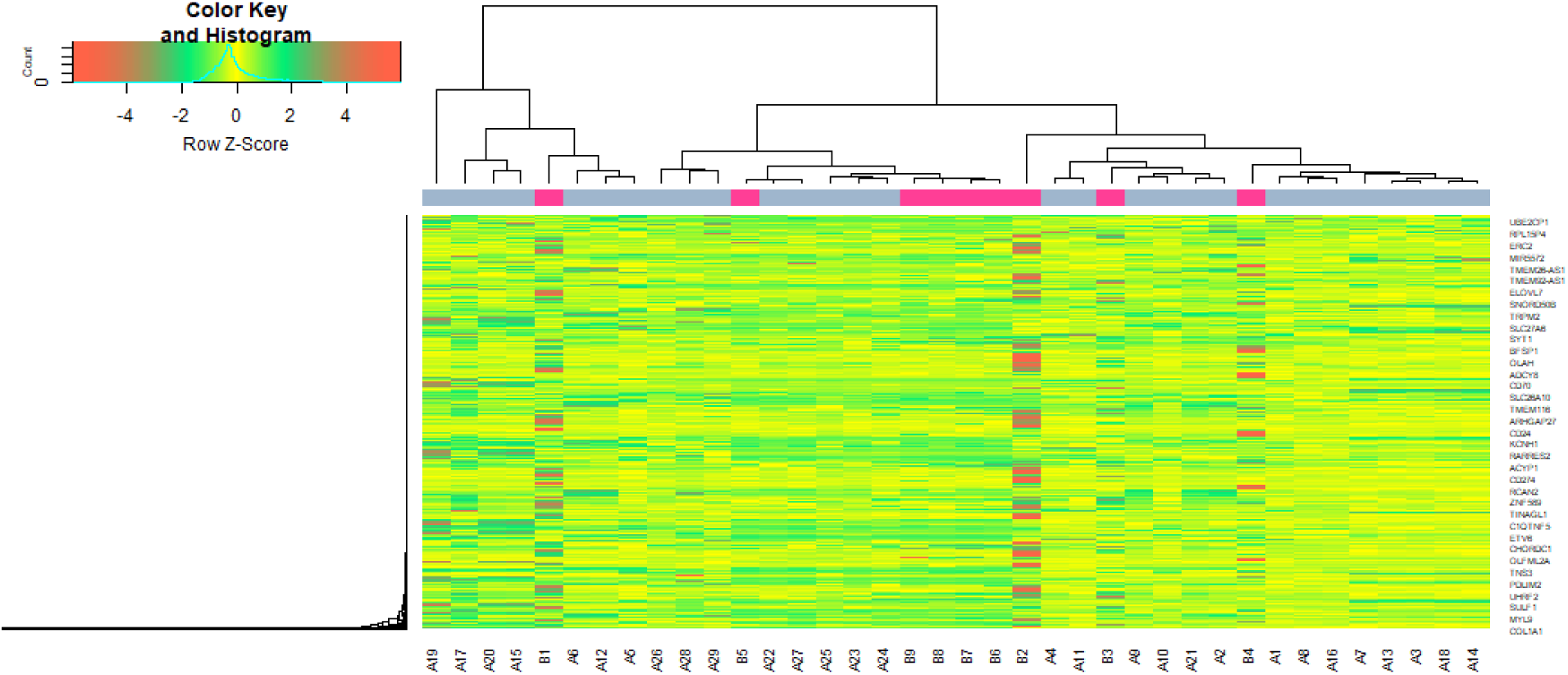
Heat map of differentially expressed genes. Legend on the top left indicate log fold change of genes. (A1 – A29 = DKD samples; B1 – B9 = normal control samples)

### GO and pathway enrichment analyses of DEGs

A total of 479 up regulated genes and 479 down regulated genes were analyzed by g:Profiler software. The significant terms from the GO enrichment analysis showed that in the BP category, the up regulated genes were involved in biological regulation and cellular response to stimulus, whereas the down regulated genes were significantly involved in the multicellular organismal process and cell communication (Table 2). For the CC category, the up regulated genes were correlated with the cell periphery and membrane, whereas the down regulated genes were associated with the cytoplasm and vesicle (Table 2). For the MF category, the up regulated genes were enriched in signaling receptor binding and molecular function regulator activity, whereas the down regulated genes were related to metal ion binding and cation binding (Table 2). For REACTOME pathway enrichment analysis, the up regulated genes were enriched in signaling by GPCR and GPCR ligand binding, whereas the significant REACTOME pathways of the down regulated genes included extracellular matrix organization and signal transduction (Table 3).

**Table 2.**
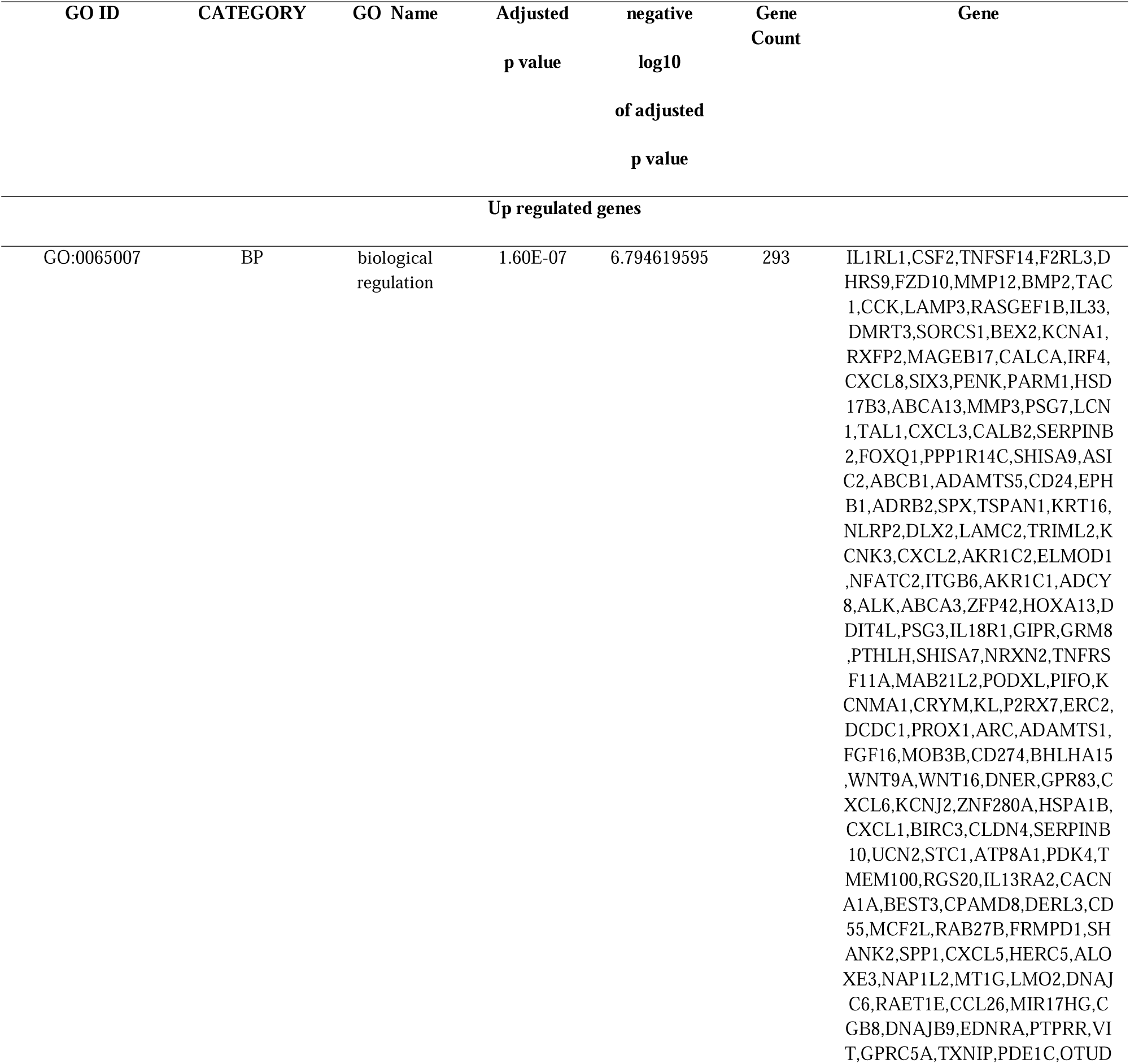
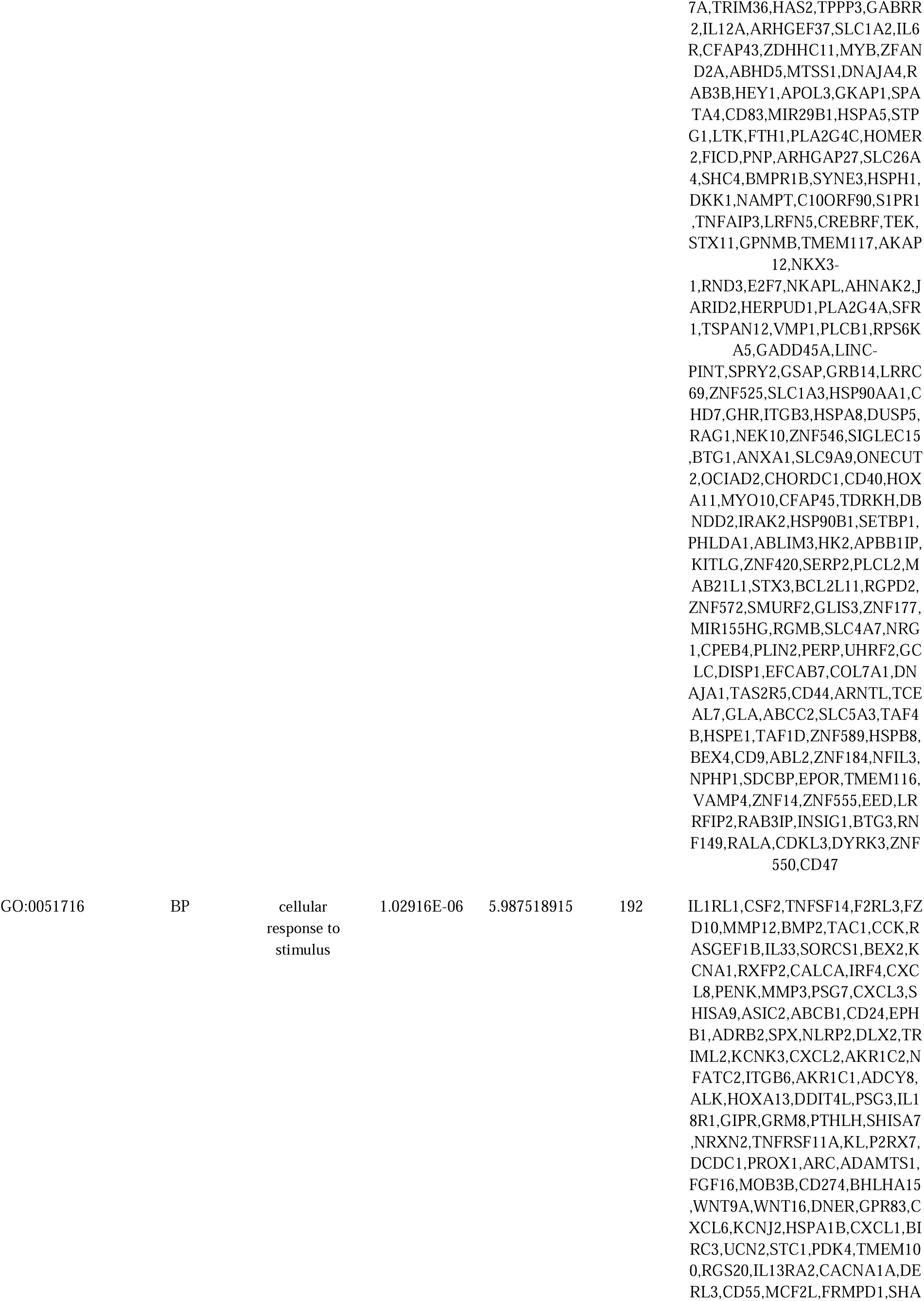
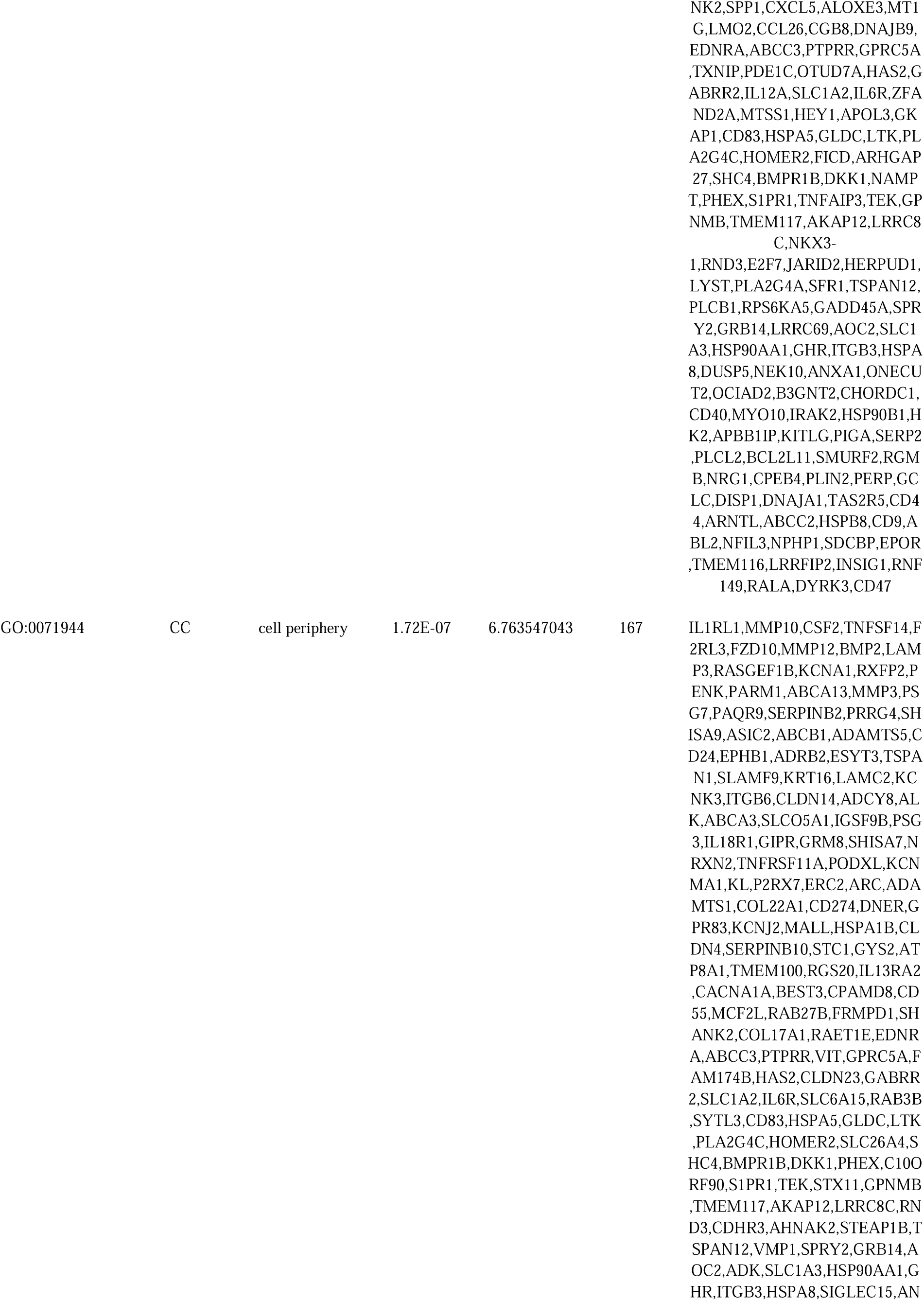
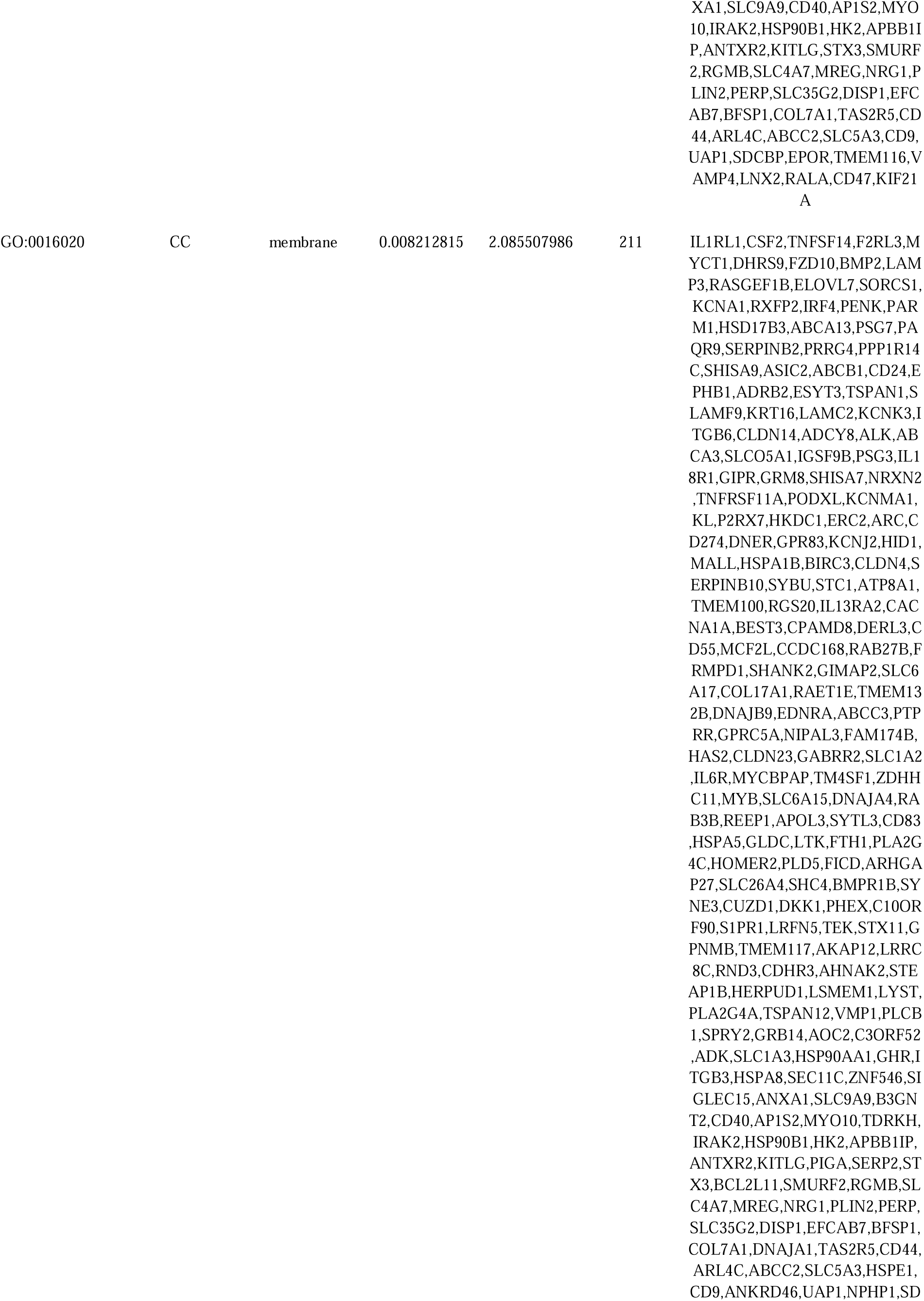
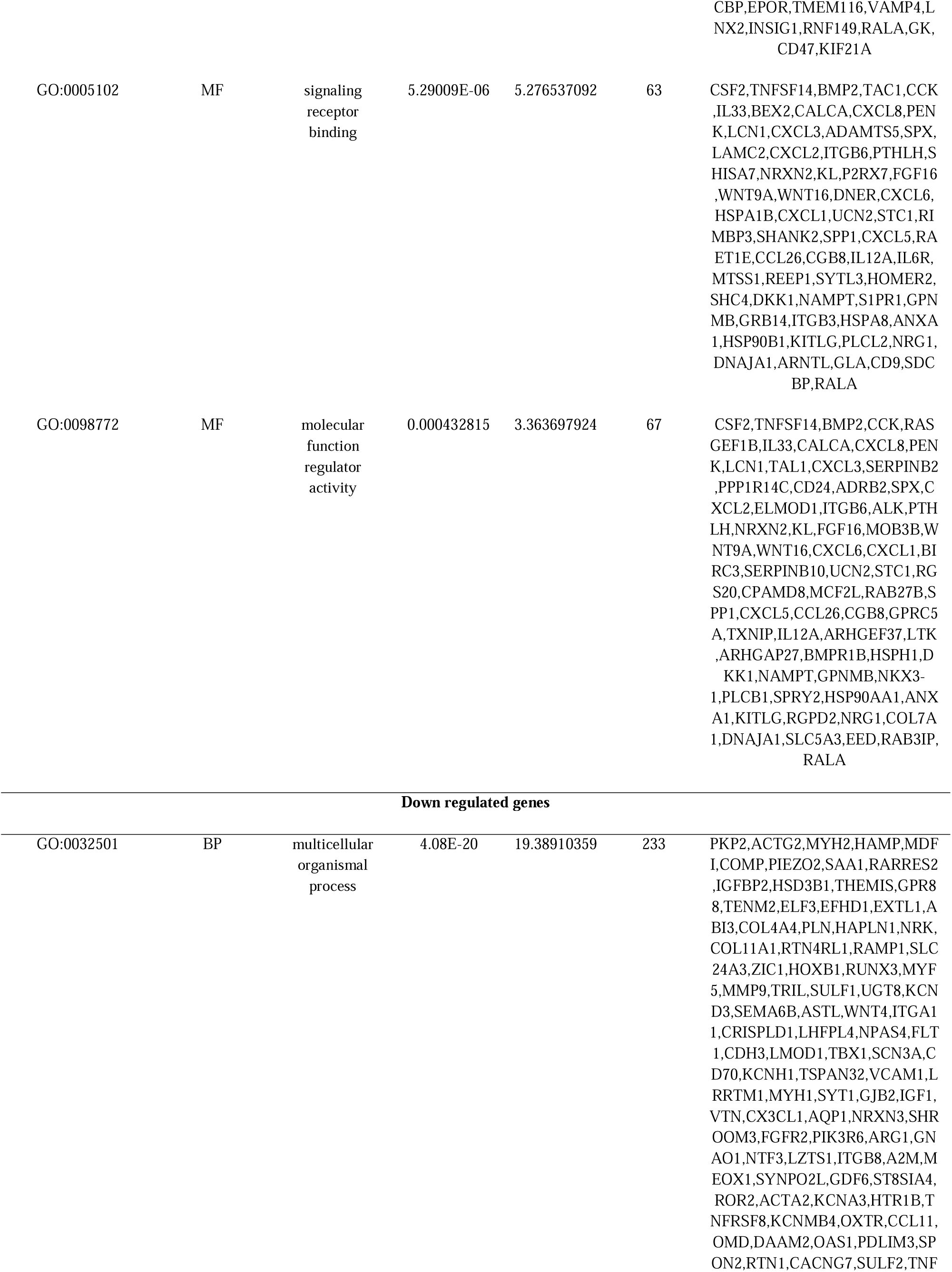
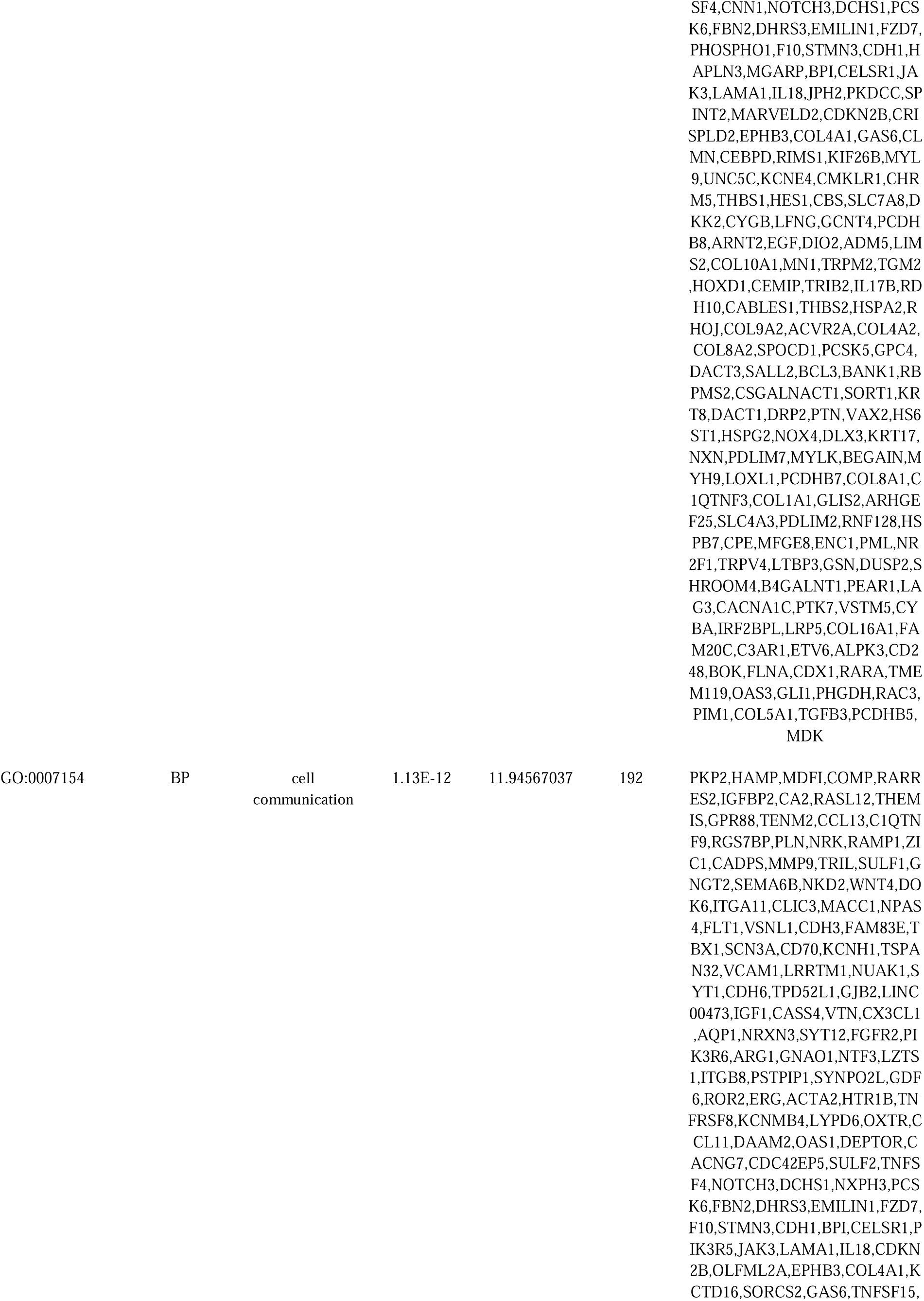
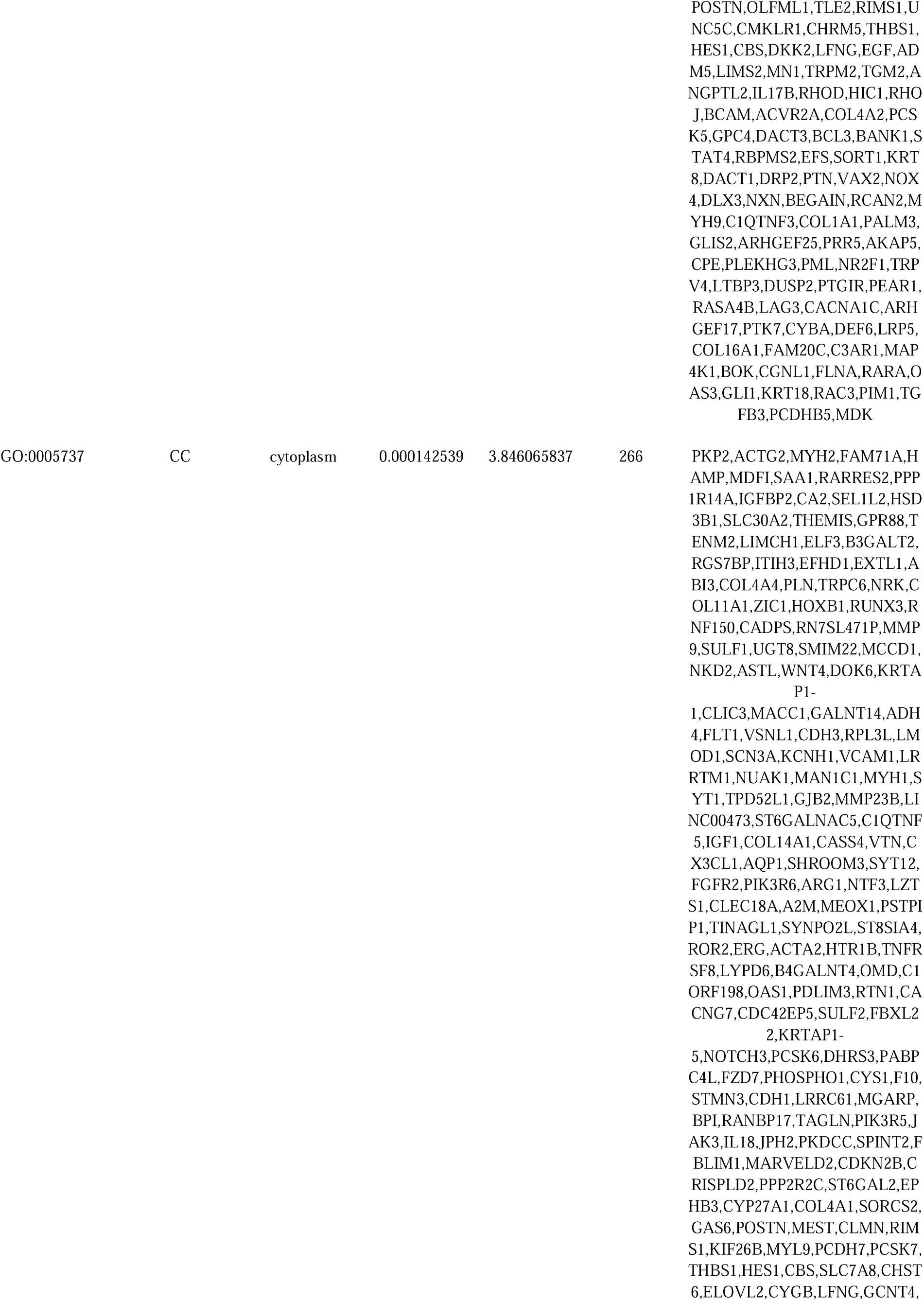
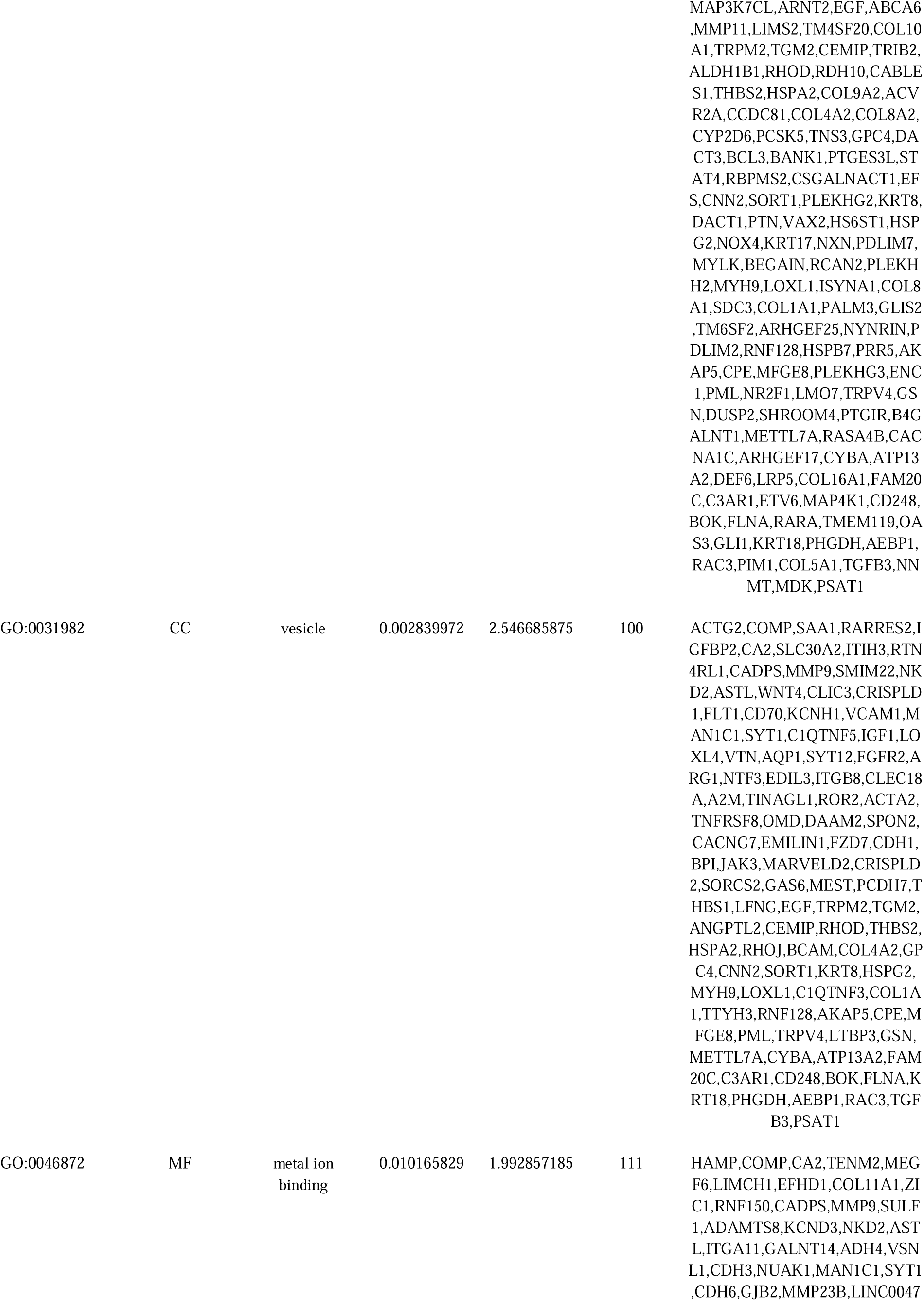
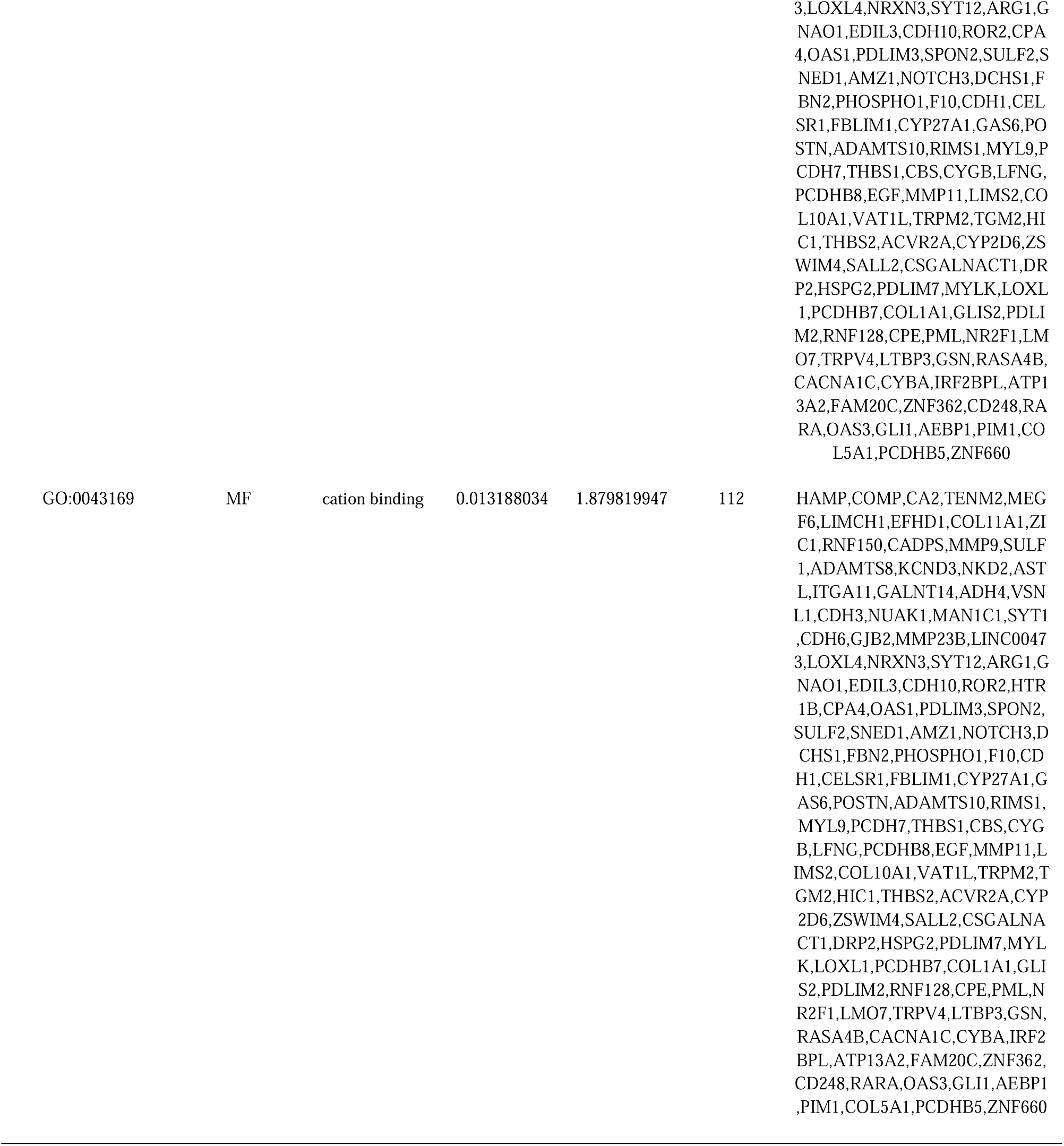
The enriched GO terms of the up and down regulated differentially expressed genes

**Table 3.**
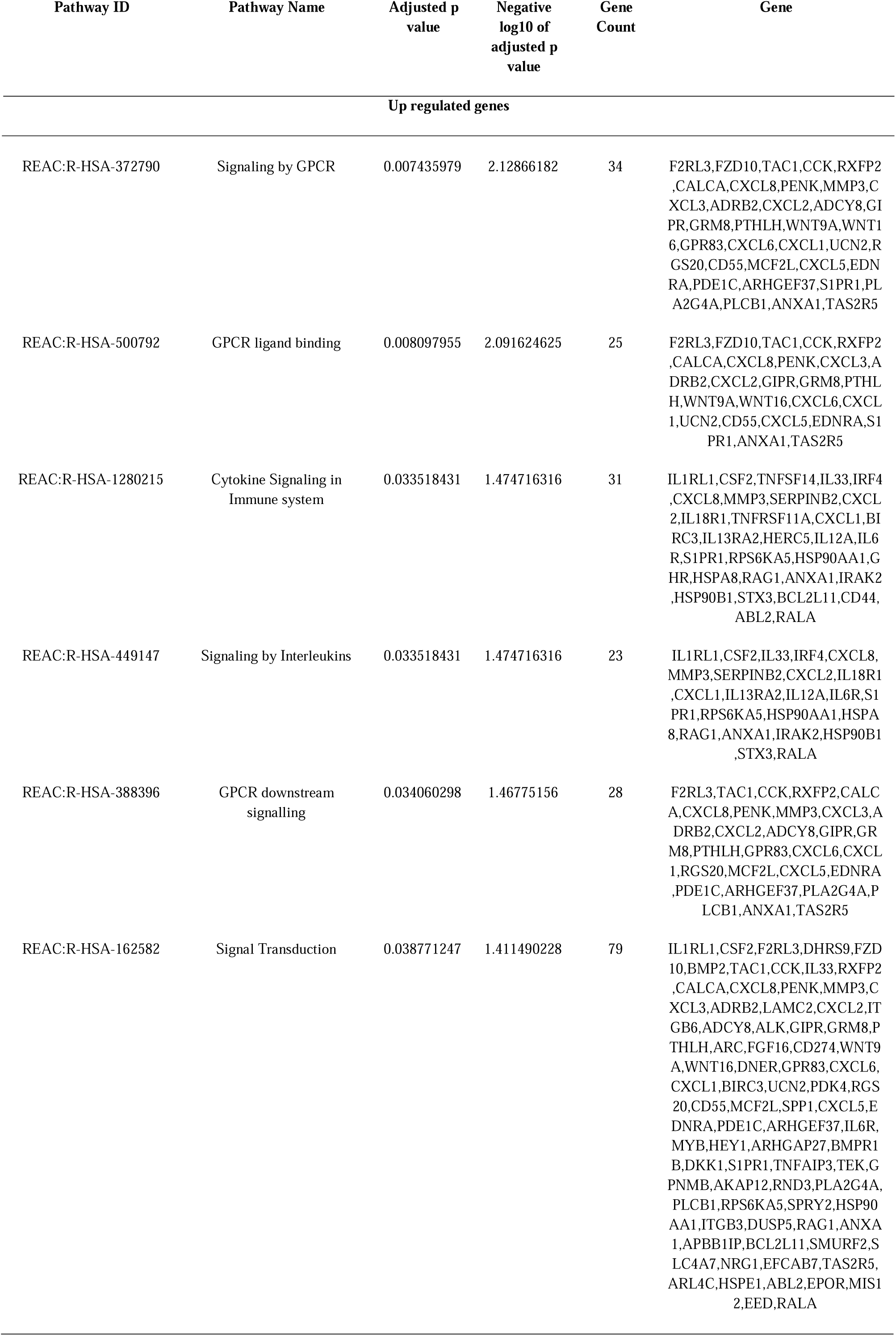
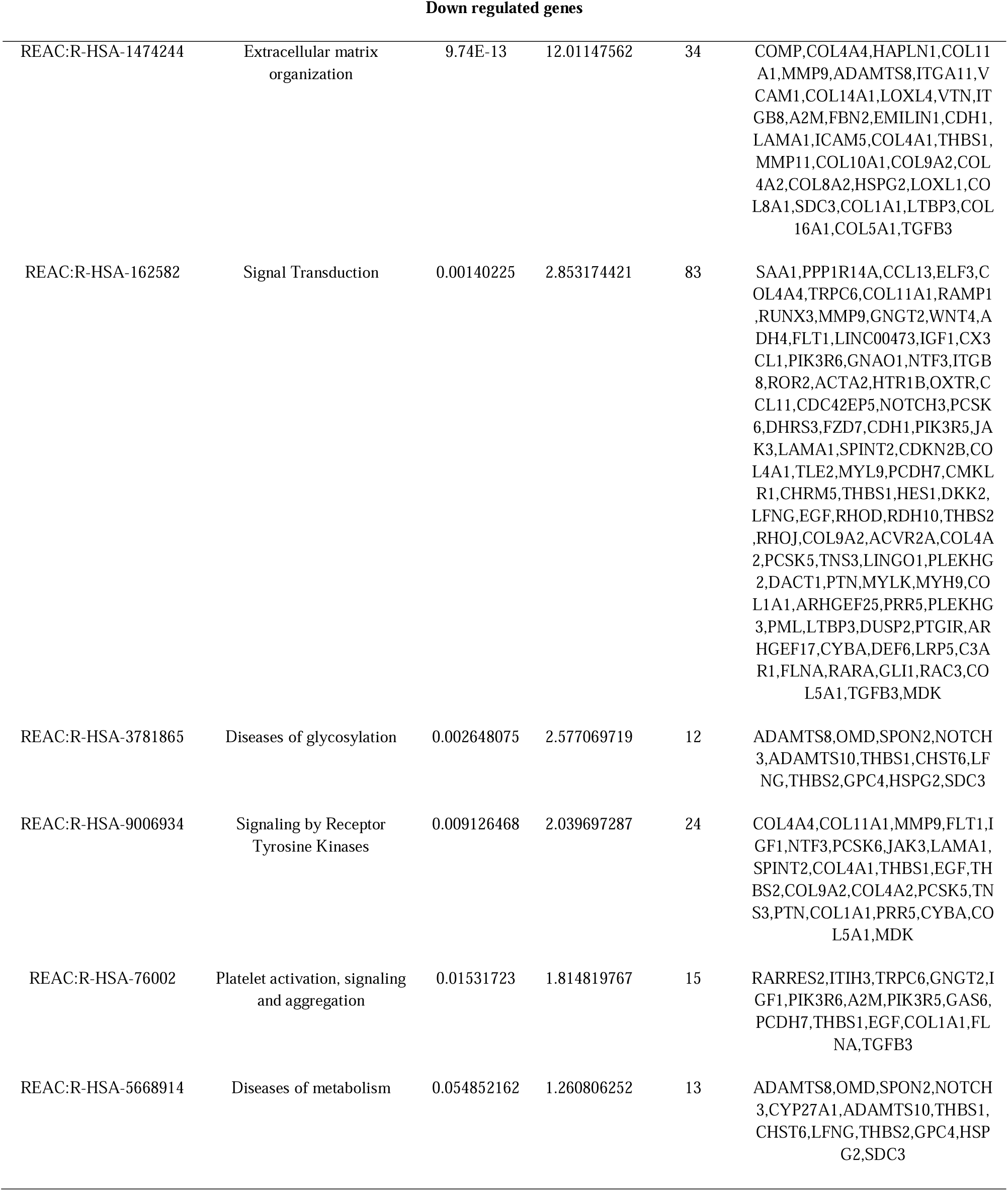
The enriched pathway terms of the up and down regulated differentially expressed genes

### Construction of the PPI network and module analysis

The PPI network of the DEGs in DKD was constructed based on the information obtained from the HIPPIE PPI database. The PPI network included 5516 nodes and 9839 edges (Fig. 3). HSPA8, HSP90AA1, HSPA5, SDCBP, HSP90B1, VCAM1, MYH9, FLNA, MDFI and PML are identified as hub genes associated with DKD (Table 4). Then, the significant modules were identified via the PEWCC1 plugin. The top two functional clusters of modules were selected. GO and REACTOME enrichment pathway analysis of each module was performed by g:Profiler. Module 1 consisted of 89 nodes and 240 edges (Fig. 4A) with genes enriched in cytokine signaling in immune system, signaling by interleukins and biological regulation. Module 2 consisted of 59 nodes and 157 edges (Fig. 4B) with genes enriched in signal transduction and multicellular organismal process.

**Fig. 3.**
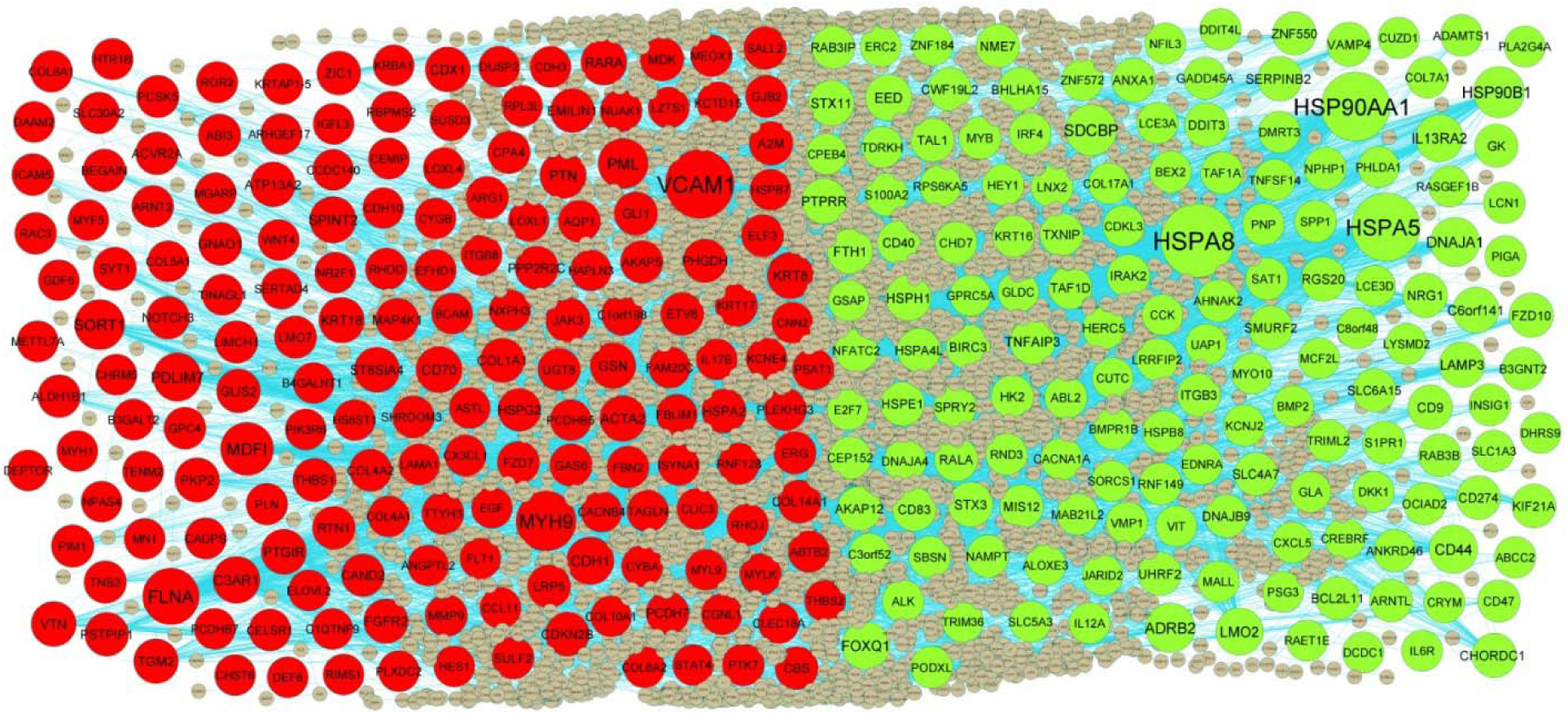
PPI network of DEGs. Up regulated genes are marked in parrot green; down regulated genes are marked in red

**Fig. 4.**
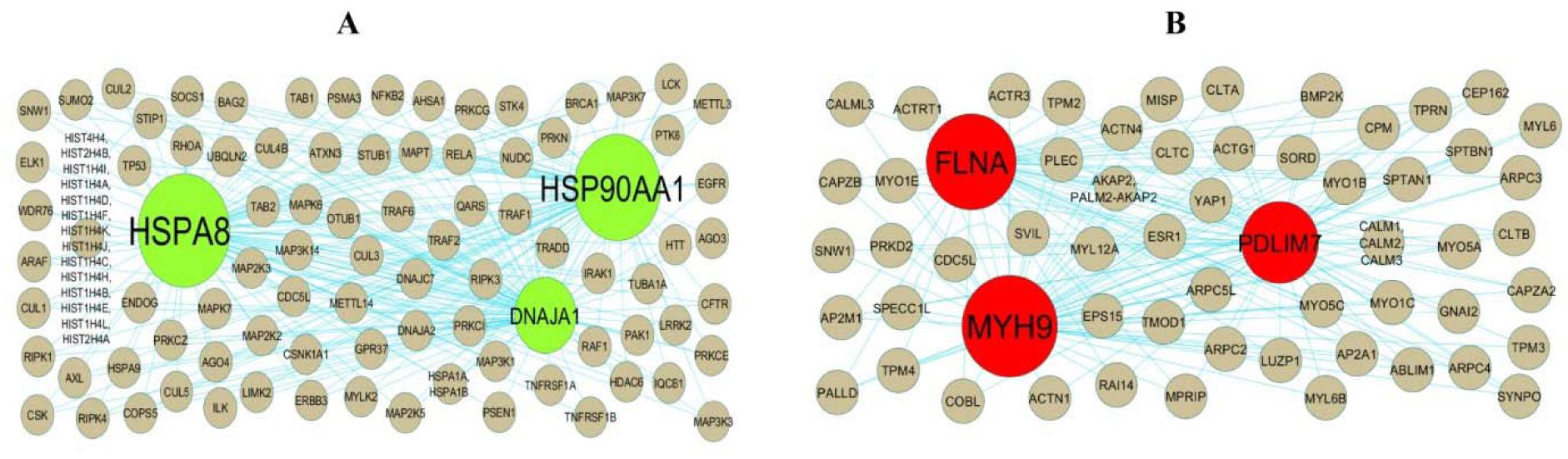
Modules selected from the DEG PPI between patients with FSGS and normal controls. (A) The most significant module was obtained from PPI network with 89 nodes and 240 edges for up regulated genes (B) The most significant module was obtained from PPI network with 59 nodes and 157 edges for down regulated genes. Up regulated genes are marked in parrot green; down regulated genes are marked in red

**Table 4.**
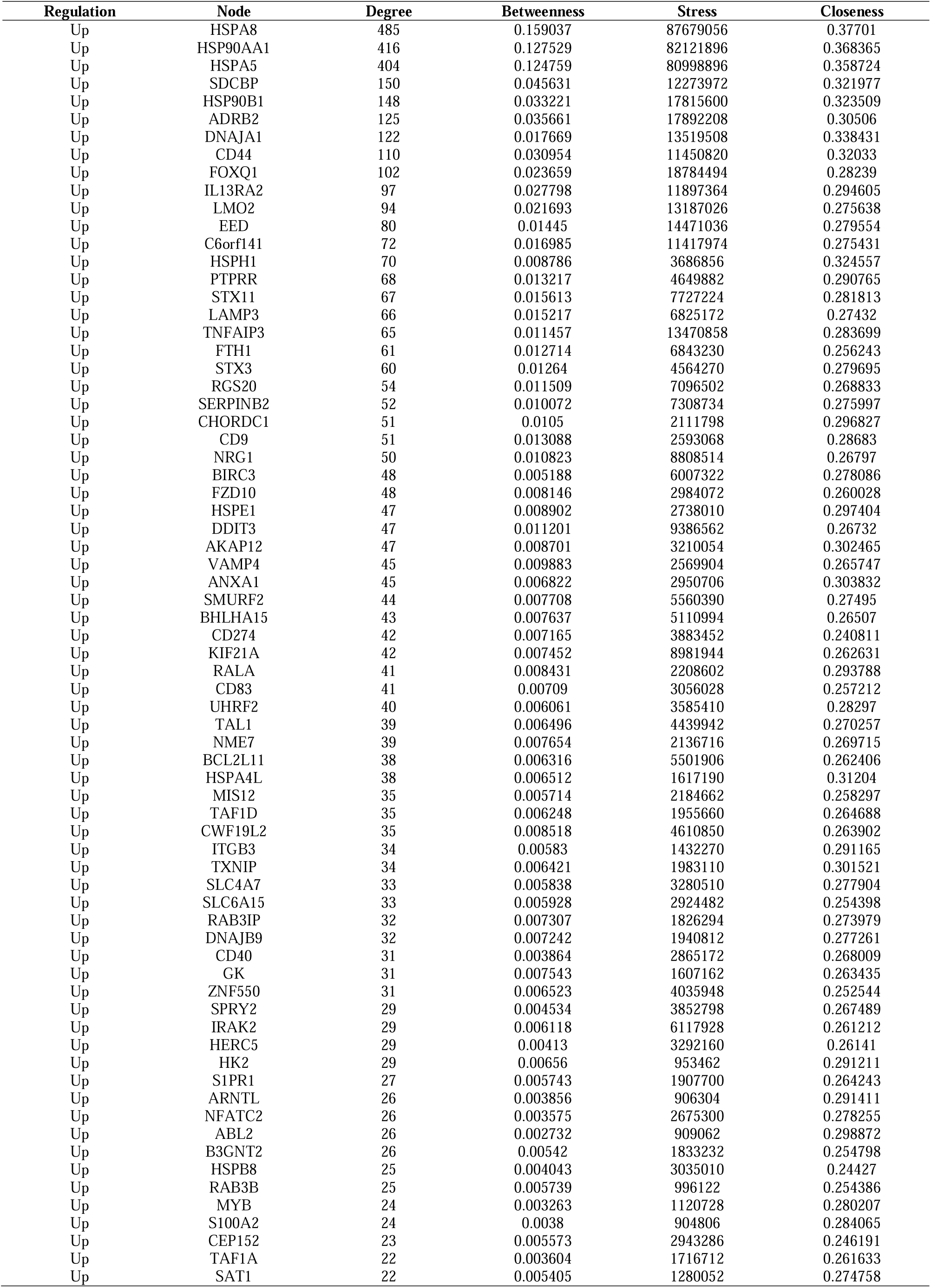
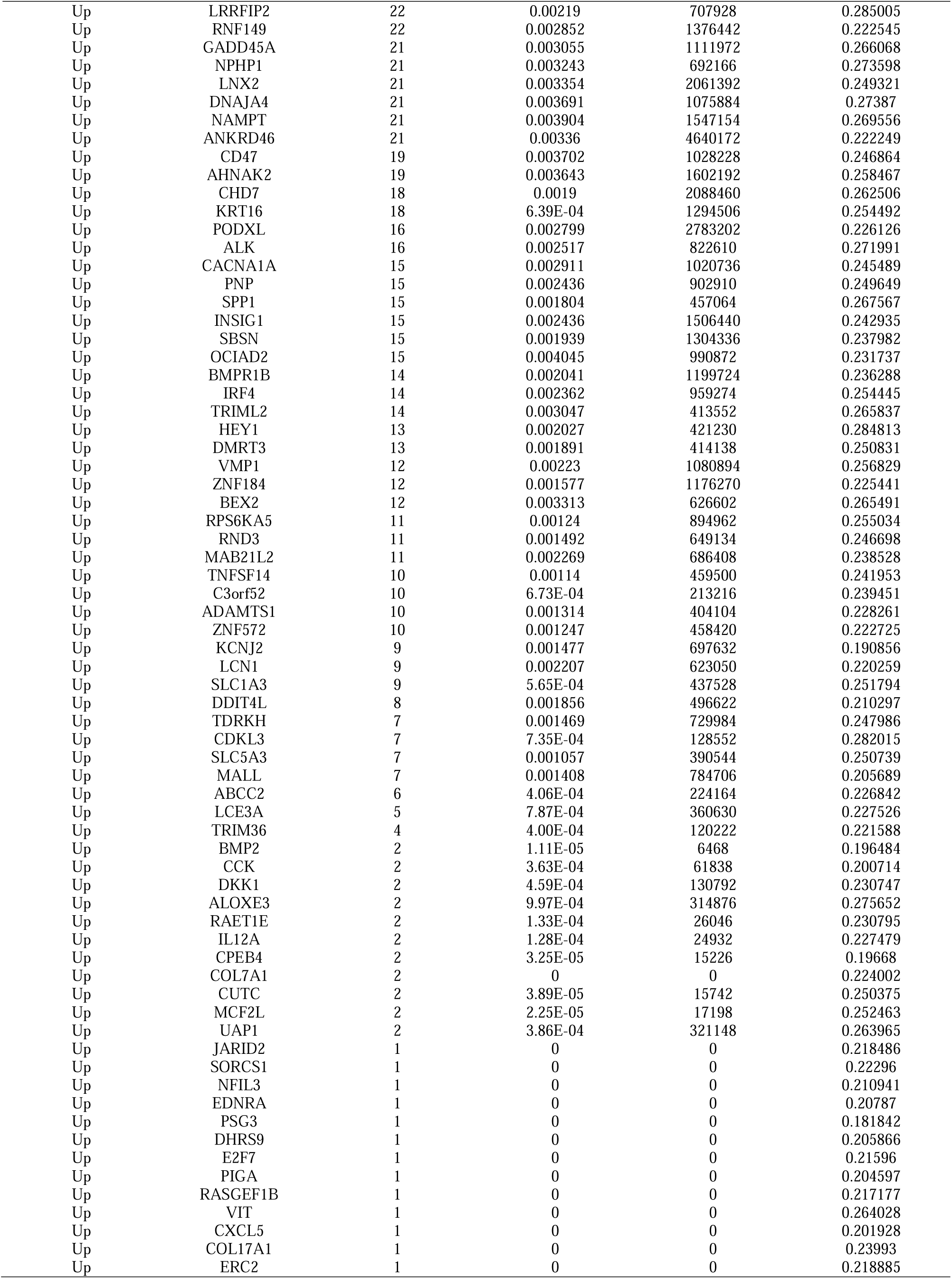
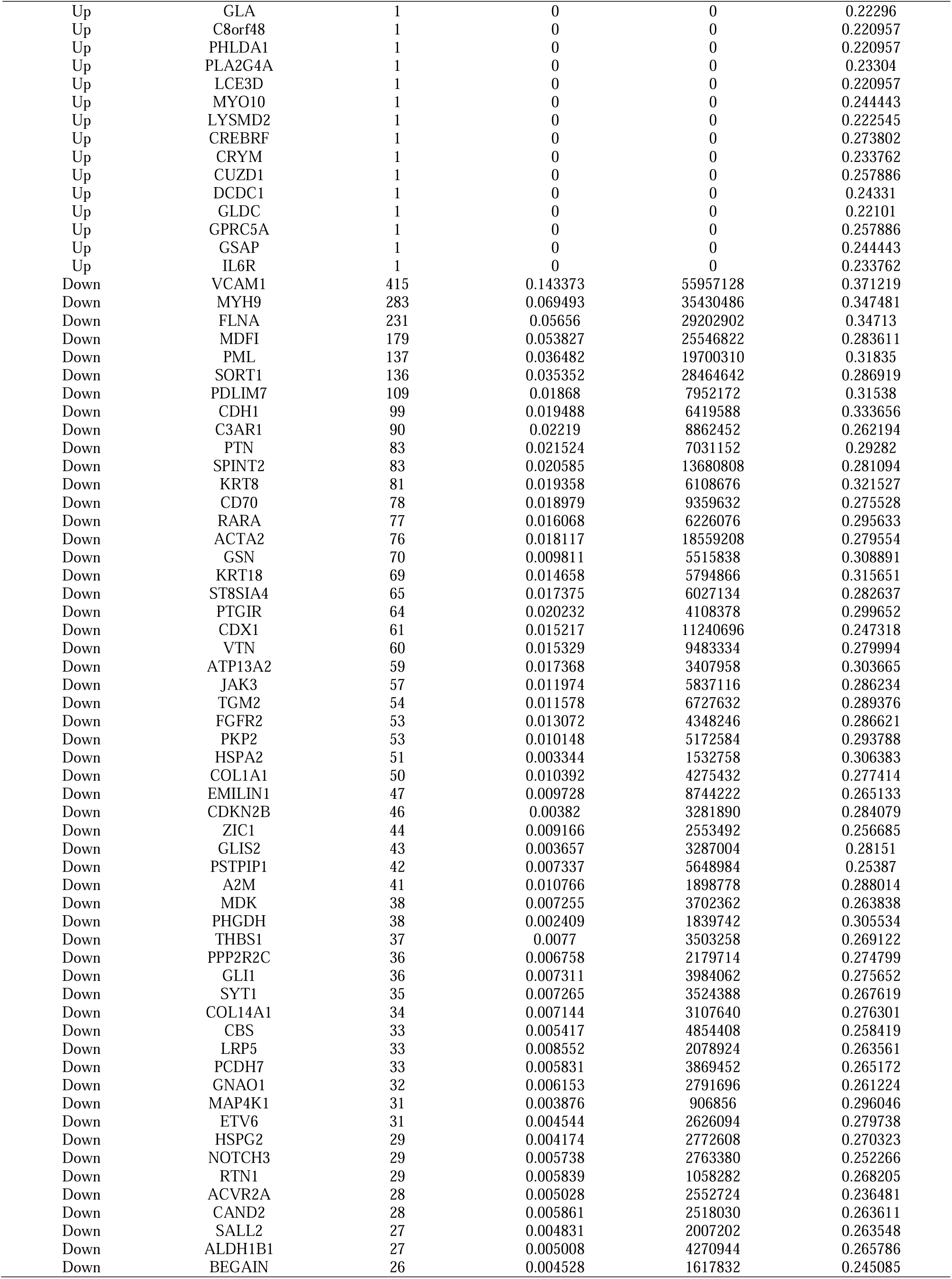
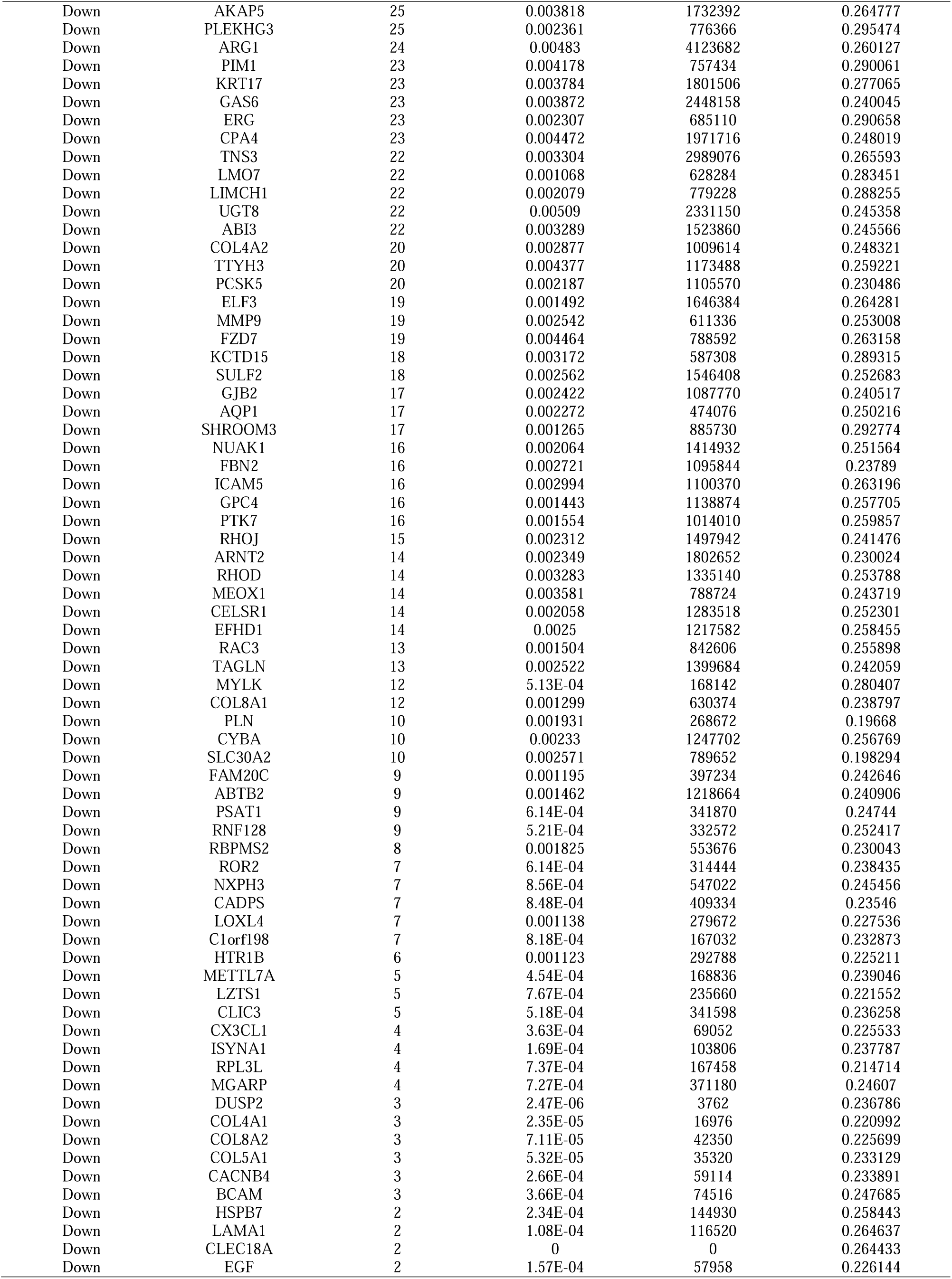
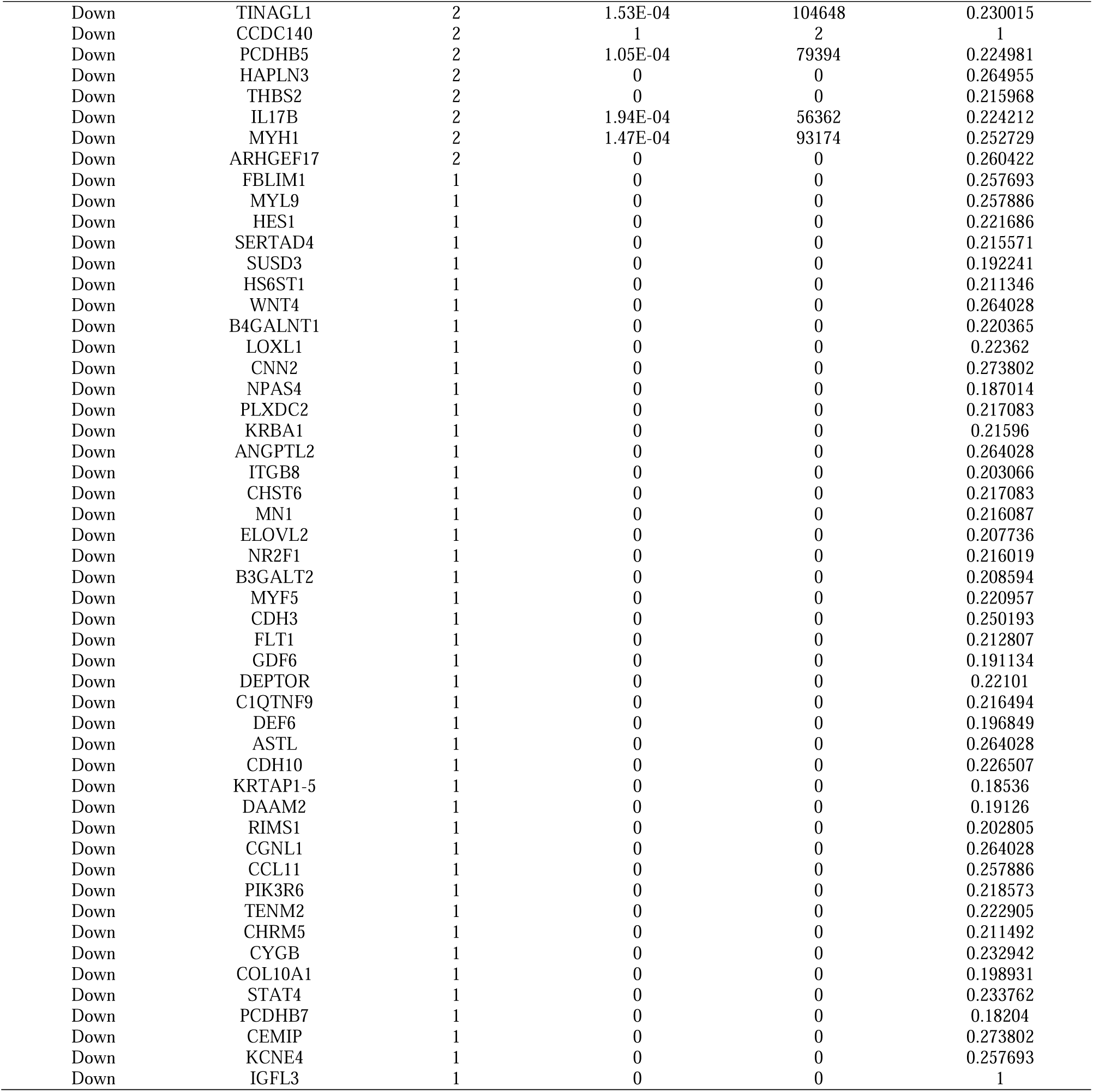
Topology table for up and down regulated genes

### Construction of the miRNA-hub gene regulatory network

The miRNet database and Cytoscape software were used to establish the miRNA-hub gene regulatory network of the hub genes. A miRNA-hub gene regulatory network containing 2590 nodes (miRNAs: 2259; Hub Genes: 331) and 16521 edges was constructed (Fig. 5). HSPA8 that was modulated by 116 miRNAs (ex: hsa-mir-606); HSP90AA1 that was modulated by 106 miRNAs (ex: hsa-mir-301a-3p); HSP90B1 that was modulated by 104 miRNAs (ex: hsa-mir-129-1-3p); HSPH1 that was modulated by 91 miRNAs (ex: hsa-mir-152-3p); HSPA5 that was modulated by 88 miRNAs (ex: hsa-mir-5006-3p); MYH9 that was modulated by 126 miRNAs (ex: hsa-mir-412-3p); FLNA that was modulated by 92 miRNAs (ex: hsa-mir-653-5p); CDH1 that was modulated by 60 miRNAs (ex: hsa-mir-647); SORT1 that was modulated by 57 miRNAs (ex: hsa-mir-125a-5p); ACTA2 that was modulated by 40 miRNAs (ex: hsa-mir-375) (Table 5).

**Fig. 5.**
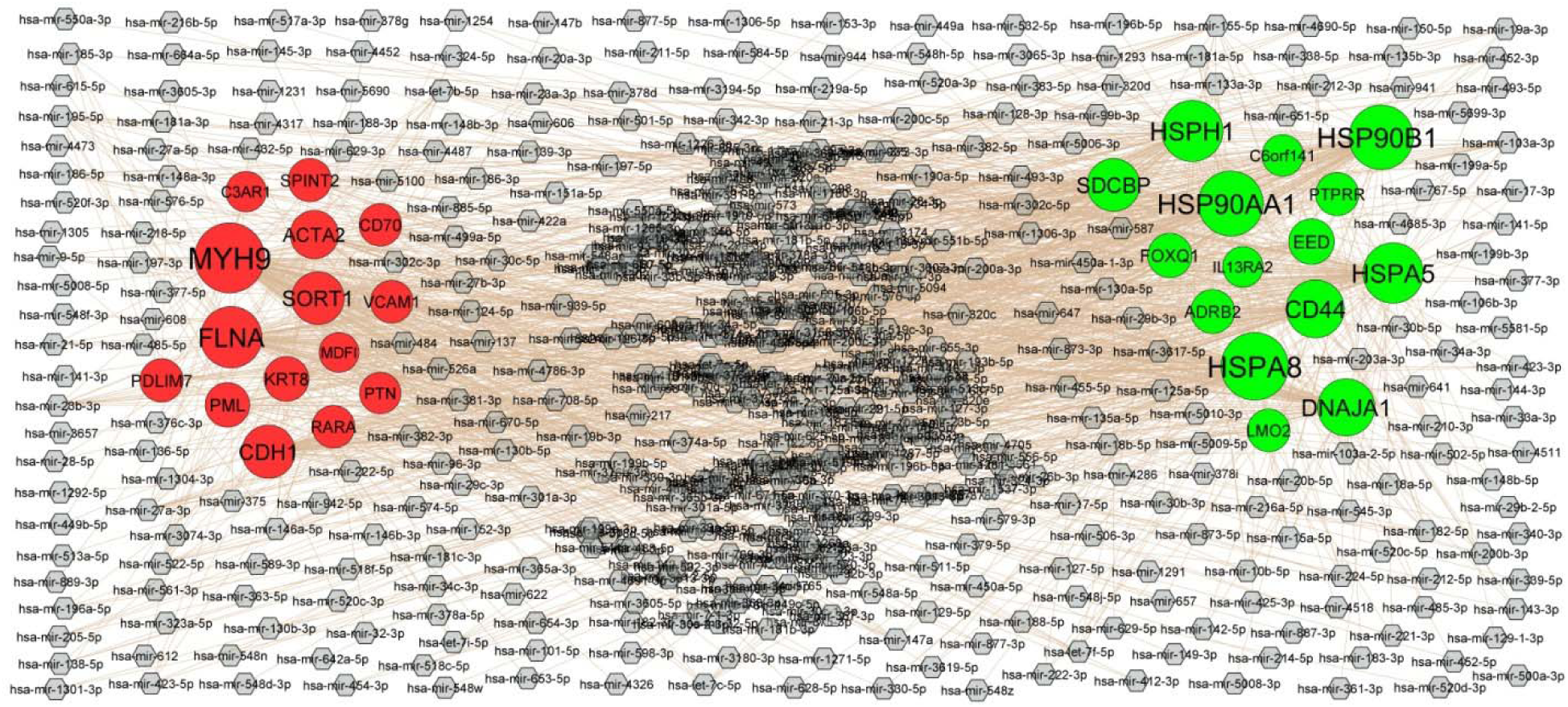
Target gene - miRNA regulatory network between target genes. The ash color diamond nodes represent the key miRNAs; up regulated genes are marked in green; down regulated genes are marked in red.

**Table 5.** MiRNA - hub gene and TF – hub gene topology table

| Regulation | Target Genes | Degree | MicroRNA | Regulation | Target Genes | Degree | TF |
| --- | --- | --- | --- | --- | --- | --- | --- |
| Up | HSPA8 | 116 | hsa-mir-606 | Up | HSP90AA1 | 45 | SOX2 |
| Up | HSP90AA1 | 106 | hsa-mir-301a-3p | Up | LMO2 | 45 | SMAD4 |
| Up | HSP90B1 | 104 | hsa-mir-129-1-3p | Up | HSPA5 | 44 | GATA2 |
| Up | HSPH1 | 91 | hsa-mir-152-3p | Up | EED | 41 | CCND1 |
| Up | HSPA5 | 88 | hsa-mir-5006-3p | Up | HSPA8 | 39 | CREM |
| Up | CD44 | 78 | hsa-mir-526a | Up | STX11 | 34 | PAX3 |
| Up | DNAJA1 | 75 | hsa-mir-15a-5p | Up | FOXQ1 | 34 | MYC |
| Up | SDCBP | 57 | hsa-mir-203a-3p | Up | HSP90B1 | 31 | PADI4 |
| Up | EED | 29 | hsa-mir-30c-5p | Up | CD44 | 31 | BACH1 |
| Up | FOXQ1 | 24 | hsa-mir-148b-3p | Up | SDCBP | 29 | ESR2 |
| Up | ADRB2 | 22 | hsa-mir-17-5p | Up | ADRB2 | 27 | CNOT3 |
| Up | PTPRR | 19 | hsa-mir-27a-5p | Up | DNAJA1 | 27 | TTF2 |
| Up | LMO2 | 17 | hsa-mir-4511 | Up | PTPRR | 24 | SALL4 |
| Up | C6orf141 | 13 | hsa-mir-1306-5p | Up | HSPH1 | 24 | MYB |
| Up | IL13RA2 | 9 | hsa-mir-374a-5p | Up | IL13RA2 | 8 | RUNX2 |
| Down | MYH9 | 126 | hsa-mir-412-3p | Down | MYH9 | 53 | EP300 |
| Down | FLNA | 92 | hsa-mir-653-5p | Down | RARA | 42 | SPI1 |
| Down | CDH1 | 60 | hsa-mir-647 | Down | CDH1 | 37 | SMARCA4 |
| Down | SORT1 | 57 | hsa-mir-125a-5p | Down | PML | 32 | HOXC9 |
| Down | ACTA2 | 40 | hsa-mir-375 | Down | KRT8 | 31 | TCF3 |
| Down | KRT8 | 26 | hsa-mir-378a-5p | Down | PTN | 27 | WT1 |
| Down | PML | 25 | hsa-mir-19b-3p | Down | SPINT2 | 26 | GATA3 |
| Down | PDLIM7 | 20 | hsa-mir-23b-3p | Down | SORT1 | 19 | POU5F1 |
| Down | SPINT2 | 20 | hsa-mir-493-5p | Down | FLNA | 18 | DMRT1 |
| Down | RARA | 17 | hsa-mir-1291 | Down | PDLIM7 | 18 | RARG |
| Down | VCAM1 | 17 | hsa-mir-195-5p | Down | MDF1 | 17 | SUZ12 |
| Down | CD70 | 17 | hsa-mir-147a | Down | VCAM1 | 16 | RELA |
| Down | PTN | 12 | hsa-mir-449b-5p | Down | ACTA2 | 14 | JUN |
| Down | C3AR1 | 9 | hsa-mir-212-3p | Down | CD70 | 13 | SOX11 |
| Down | MDF1 | 6 | hsa-mir-103a-3p | Down | C3AR1 | 12 | ATF3 |

### Construction of the TF-hub gene regulatory network

The NetworkAnalyst database and Cytoscape software were used to establish the TF-hub gene regulatory network of the hub genes. A TF-hub gene regulatory network containing 520 nodes (TFs: 187; Hub Genes: 333) and 8622 edges was constructed (Fig. 6). HSP90AA1 that was modulated by 45 TFs (ex: SOX2); LMO2 that was modulated by 45 TFs (ex: SMAD4); HSPA5 that was modulated by 44 TFs (ex: GATA2); EED that was modulated by 41 TFs (ex: CCND1); HSPA8 that was modulated by 39 TFs (ex: CREM); MYH9 that was modulated by 53 TFs (ex: EP300); RARA that was modulated by 42 TFs (ex: SPI1); CDH1 that was modulated by 37 TFs (ex: SMARCA4); PML that was modulated by 32 TFs (ex: HOXC9); KRT8 that was modulated by 31 TFs (ex: TCF3) (Table 5).

**Fig. 6.**
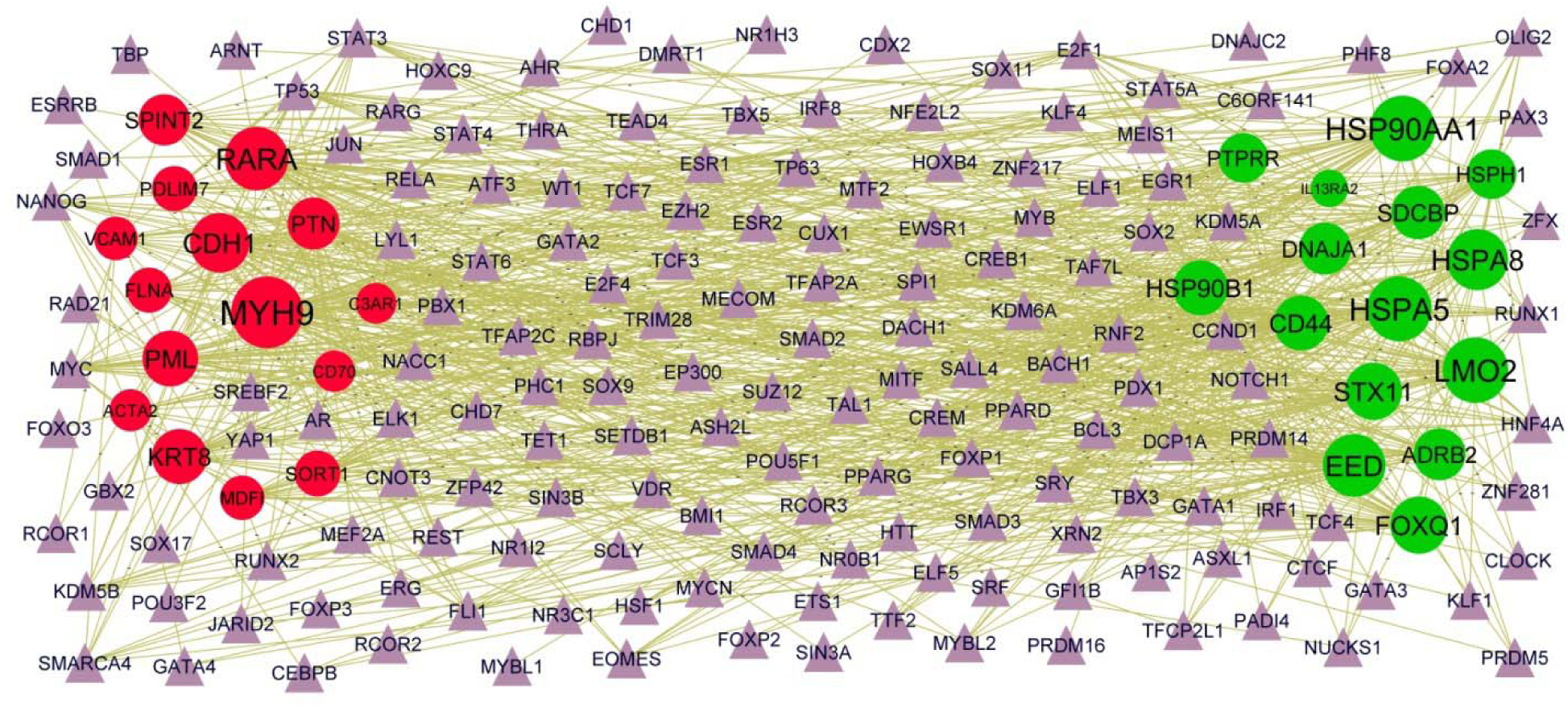
Target gene - TF regulatory network between target genes. The violet color triangle nodes represent the key TFs; up regulated genes are marked in green; down regulated genes are marked in red.

### Receiver operating characteristic curve (ROC) analysis

To determine which hub genes have the diagnose significance of DKD patients; The ROC analyses were conducted to explore the sensitivity and specificity of hub genes for DKD diagnosis. The results showed that HSPA8, HSP90AA1, HSPA5, SDCBP, HSP90B1, VCAM1, MYH9, FLNA, MDFI and PML have the best diagnostic value for differentiating the patients with DKD from normal controls. The AUC values for hub genes were HSPA8 (0.930), HSP90AA1 (0.947), HSPA5 (0.939), SDCBP (0.936), HSP90B1 (0.920), VCAM1 (0.937), MYH9 (0.944), FLNA (0.912), MDFI (0.922) and PML (0.926) (Fig. 7). This indicated that HSPA8, HSP90AA1, HSPA5, SDCBP, HSP90B1, VCAM1, MYH9, FLNA, MDFI and PML could act as biomarkers to estimate the activity of DKD and verify the effectiveness of the treatment of DKD.

**Fig. 7.**
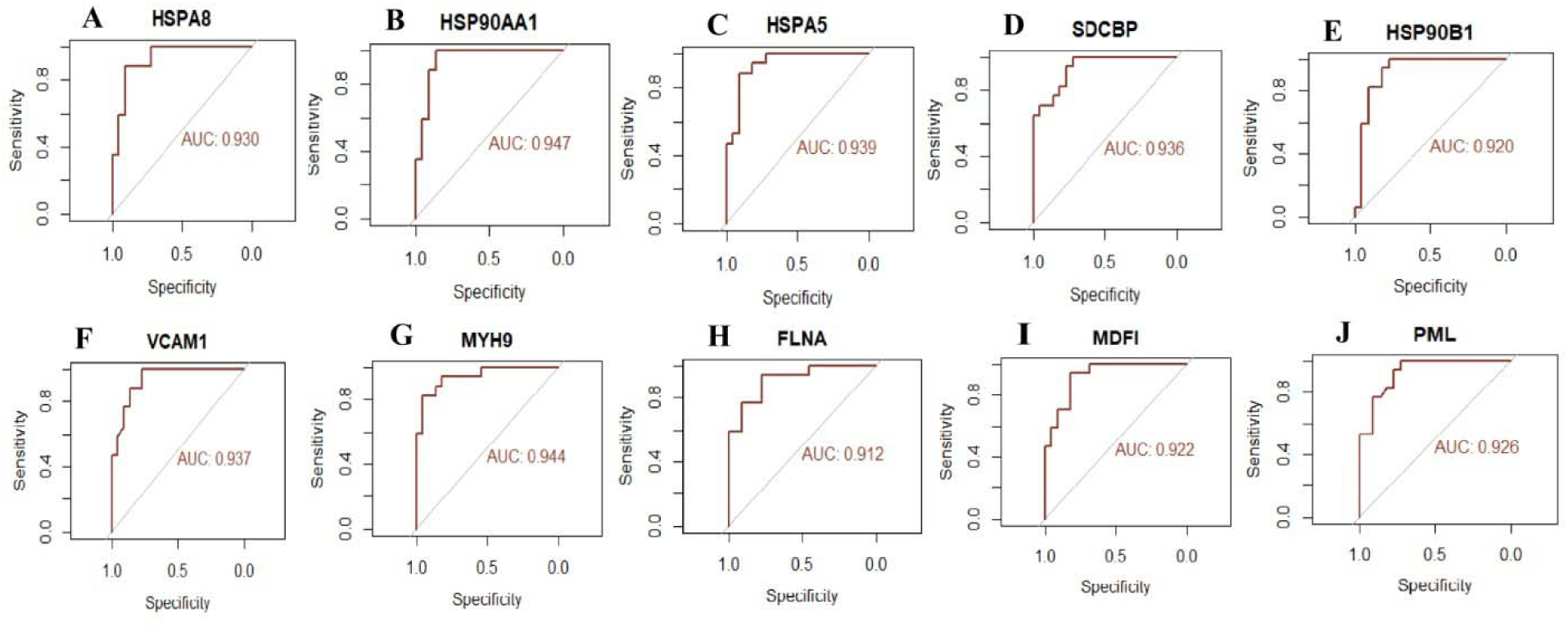
ROC curve analyses of hub genes. A) HSPA8 B) HSP90AA1 C) HSPA5 D) SDCBP E) HSP90B1 F) VCAM1 G) MYH9 H) FLNA I) MDFI J) PML

## Discussions

It is widely believed that DKD contain complex biological processes of multiple factors and stages. In current years, a large number of new biomarkers have been developed for early diagnosis, molecular pathological mechanism research, and drug target screening in the DKD [39]. Although research on DKD has increased in recent years, the molecular pathogenesis of DKD is still unclear, and the therapeutic effect is unsatisfactory.

In the current investigation, bioinformatics methods were used to analyze the NGS data of patients with DKD. In the current investigation, integrated bioinformatics methods assisted in an analysis of how key genes change in their expression to uncover potential DKD based on NGS dataset (GSE217709), and we identified total 958 DEGs, including 479 up regulated genes and 479 down regulated genes. IL1RL1 [40], MMP10 [41], TNFSF14 [42], F2RL3 [43], PKP2 [44], HAMP (hepcidin antimicrobial peptide) [45] and COMP (cartilage oligomeric matrix protein) [46] are associated with the prognosis of cardiovascular diseases. IL1RL1 [47], MMP10 [48], TNFSF14 [49], HAMP (hepcidin antimicrobial peptide) [50] and COMP (cartilage oligomeric matrix protein) [51] plays an important role in the obesity. MMP10 [52] has been identified as a key gene in DKD. MMP10 [53], TNFSF14 [54], F2RL3 [55], COMP (cartilage oligomeric matrix protein) [56] and PIEZO2 [57] are associated with the pathogenesis and development of hypertension. MMP10 [58], CSF2 [59] and HAMP (hepcidin antimicrobial peptide) [60] expression were altered in chronic kidney disease. MMP10 [61], TNFSF14 [49], HAMP (hepcidin antimicrobial peptide) [62] and COMP (cartilage oligomeric matrix protein) [63] are important in the development of type 2 diabetes mellitus. Recent studies have proposed that the MMP10 [64] and CSF2 [65] are associated with type 1 diabetes mellitus. TNFSF14 [66], HAMP (hepcidin antimicrobial peptide) [67] and COMP (cartilage oligomeric matrix protein) [68] are a pathogenic genes for renal fibrosis. These results suggested that these genes might influence the development of DKD.

Furthermore, we investigated the biological functions of these DEGs involvement in GO terms and pathways. Extracellular matrix organization [69], diseases of glycosylation [70] and diseases of metabolism [71] are responsible for progression DKD. MMP12 [72], BMP2 [73], IL33 [74], IRF4 [75], CXCL8 [76], PENK (proenkephalin) [77], SERPINB2 [78], ABCB1 [79], ADAMTS5 [80], SPX (spexin hormone) [81], HOXA13 [82], PODXL (podocalyxin like) [83], KL (klotho) [84], ADAMTS1 [85], WNT9A [86], CXCL6 [87], CXCL1 [89], STC1 [90], DNAJB9 [91], TXNIP (thioredoxin interacting protein) [92], DKK1 [93], NAMPT (nicotinamidephosphoribosyltransferase) [94], ANXA1 [95], CD40 [96], SMURF2 [97], NRG1 [98], CD44 [99], NPHP1 [100], EPOR (erythropoietin receptor) [101], CD47 [102], SAA1 [103], IGFBP2 [104], MMP9 [105], WNT4 [106], FLT1 [107], TBX1 [108], CD70 [109], VCAM1 [110], IGF1 [111], VTN (vitronectin) [112], CX3CL1 [113], AQP1 [114], SHROOM3 [115], FGFR2 [116], ROR2 [117], RTN1 [118], NOTCH3 [119], JAK3 [120], IL18 [121], COL4A1 [122], CEBPD (CCAAT enhancer binding protein delta) [123], KIF26B [124], HES1 [125], CBS (cystathionine beta-synthase) [126], EGF (epidermal growth factor) [127], TRPM2 [128], HSPG2 [129], NOX4 [130], MYH9 [131], GLIS2 [132], TRPV4 [133], GSN (gelsolin) [134], DUSP2 [135], LRP5 [136], CD248 [137], GLI1 [138], PIM1 [139], TGFB3 [140], MDK (midkine) [141], TRPC6 [142], NKD2 [143], NUAK1 [144], CYS1 [145], POSTN (periostin) [146], STAT4 [147], AEBP1 [148] and NNMT (nicotinamide N-methyltransferase) [149] might be a possible genetic markers for susceptibility to renal fibrosis. MMP12 [150], BMP2 [151], IL33 [152], SORCS1 [153], IRF4 [154], PENK (proenkephalin) [77], ABCB1 [155], ADRB2 [156], SPX (spexin hormone) [157], PODXL (podocalyxin like) [158], KL (klotho) [159], ADAMTS1 [160], CXCL1 [161], PDK4 [162], CD55 [163], MIR17HG [164], DNAJB9 [165], GPRC5A [166], TXNIP (thioredoxin interacting protein) [167], APOL3 [168], DKK1 [169], GPNMB (glycoprotein nmb) [170], GADD45A [171], GSAP (gamma-secretase activating protein) [172], GHR (growth hormone receptor) [173], HSPA8 [174], DUSP5 [175], RAG1 [176], CD40 [177], HK2 [178], GLIS3 [179], CD44 [180], NPHP1 [181], EPOR (erythropoietin receptor) [182], CD47 [183], IGFBP2 [184], COL4A4 [185], MMP9 [186], WNT4 [187], FLT1 [188], IGF1 [189], VTN (vitronectin) [190], CX3CL1 [191], AQP1 [192], SHROOM3 [193], FGFR2 [194], SPON2 [195], NOTCH3 [196], IL18 [197], COL4A1 [198], GAS6 [199], CEBPD (CCAAT enhancer binding protein delta) [200], THBS1 [201], CYGB (cytoglobin) [202], EGF (epidermal growth factor) [203], TRPM2 [204], TGM2 [205], THBS2 [174], COL4A2 [206], PCSK5 [207], NOX4 [208], MYH9 [209], GLIS2 [210], MFGE8 [211], CD248 [212], GLI1 [138], MDK (midkine) [213], TRPC6 [214], POSTN (periostin) [215], MMP11 [216], CYP2D6 [217], STAT4 [218] and NNMT (nicotinamide N-methyltransferase) [219] might be genetic factors contributing to the pathogenesis of chronic kidney disease. Altered expression of MMP12 [150], BMP2 [220], CCK (cholecystokinin) [221], IL33 [222], IRF4 [223], CXCL8 [224], PENK (proenkephalin) [225], MMP3 [226], ABCB1 [155], ADAMTS5 [227], CD24 [228], ADRB2 [229], CXCL2 [230], NFATC2 [231], DDIT4L [232], IL18R1 [233], PODXL (podocalyxin like) [234], KCNMA1 [235], KL (klotho) [236], PROX1 [237], FGF16 [238], HSPA1B [239], CXCL1 [240], UCN2 [241], PDK4 [242], CD55 [243], MCF2L [244], CXCL5 [245], RAET1E [246], TXNIP (thioredoxin interacting protein) [247], HAS2 [248], IL12A [249], IL6R [250], SLC26A4 [251], DKK1 [252], NAMPT (nicotinamidephosphoribosyltransferase) [253], TNFAIP3 [254], GPNMB (glycoprotein nmb) [255], RND3 [256], GADD45A [257], LINC-PINT [258], HSP90AA1 [259], CHD7 [260], ITGB3 [261], HSPA8 [262], DUSP5 [263], BTG1 [264], ANXA1 [95], CD40 [265], HOXA11 [266], HK2 [267], PLCL2 [268], BCL2L11 [269], MIR155HG [270], NRG1 [271], GCLC (glutamate-cysteine ligase catalytic subunit) [272], CD44 [273], HSPB8 [274], CD9 [275], EPOR (erythropoietin receptor) [276], CD47 [277], ESYT3 [278], SYTL3 [279], CDHR3 [280], ADK (adenosine kinase) [281], GPNMB (glycoprotein nmb) [282], SAA1 [283]. IGFBP2 [284], EFHD1 [285], PLN (phospholamban) [286], RUNX3 [287], MMP9 [288], KCND3 [289], WNT4 [290], CRISPLD1 [291], NPAS4 [292], FLT1 [293], LMOD1 [294], TBX1 [295], CD70 [296], VCAM1 [297], IGF1 [298], CX3CL1 [299], AQP1 [300], SHROOM3 [301], FGFR2 [302], A2M [303], MEOX1 [304], SYNPO2L [305], ROR2 [306], ACTA2 [307], CCL11 [308], OMD (osteomodulin) [309], PDLIM3 [310], SULF2 [311], TNFSF4 [312], NOTCH3 [313], PCSK6 [314], BPI (bactericidal permeability increasing protein) [315], CELSR1 [316], JAK3 [317], IL18 [318], JPH2 [319], CDKN2B [320], CRISPLD2 [321], GAS6 [322], KCNE4 [323], CMKLR1 [324], THBS1 [325], HES1 [326], CBS (cystathionine beta-synthase) [327], DKK2 [328], EGF (epidermal growth factor) [329], LIMS2 [330], TRPM2 [331], THBS2 [332], COL4A2 [320], GPC4 [333], BCL3 [334], SORT1 [335], DACT1 [336], HSPG2 [337], NOX4 [338], PDLIM7 [305], MYH9 [339], COL1A1 [340], HSPB7 [341], MFGE8 [342], TRPV4 [343], GSN (gelsolin) [344], PEAR1 [345], LAG3 [346], CACNA1C [347], CYBA (cytochrome b-245 alpha chain) [348], LRP5 [349], FAM20C [350], ALPK3 [351], GLI1 [352], PIM1 [353], TGFB3 [354], MDK (midkine) [355], ITIH3 [356], TRPC6 [357], RPL3L [358], ST6GALNAC5 [359], TAGLN (transgelin) [360], CYP27A1 [361], POSTN (periostin) [362], CYP2D6 [363], STAT4 [364], TM6SF2 [365], AKAP5 [366], AEBP1 [367] and ADAMTS8 [368] were correlated with cardiovascular diseases progression and prognosis. MMP12 [369], BMP2 [370], CCK (cholecystokinin) [371], IL33 [372], CALCA (calcitonin related polypeptide alpha) [373], CXCL8 [374], MMP3 [375], ASIC2 [376], ABCB1 [377], SPX (spexin hormone) [378], KCNK3 [379], ALK (ALK receptor tyrosine kinase) [380], ABCA3 [381], KCNMA1 [235], KL (klotho) [382], CXCL1 [383], CLDN4 [384], UCN2 [241], PDK4 [385], CACNA1A [386], CXCL5 [387], EDNRA (endothelin receptor type A) [388], TXNIP (thioredoxin interacting protein) [389], HAS2 [390], IL6R [391], SLC26A4 [392], BMPR1B [393], DKK1 [394], NAMPT (nicotinamidephosphoribosyltransferase) [395], S1PR1 [396], TNFAIP3 [397], GHR (growth hormone receptor) [398], ITGB3 [399], HSPA8 [174], DUSP5 [175], RAG1 [176], CD40 [400], HK2 [401], SLC4A7 [402], NRG1 [403], CD44 [404], ARNTL (aryl hydrocarbon receptor nuclear translocator like) [405], ABCC2 [406], EPOR (erythropoietin receptor) [407], CD47 [408], ANTXR2 [409], IGFBP2 [410], HSD3B1 [411], GPR88 [412], RAMP1 [413], MMP9 [414], WNT4 [415], FLT1 [416], CD70 [109], VCAM1 [391], IGF1 [417], CX3CL1 [418], AQP1 [419], FGFR2 [420], ARG1 [421], ACTA2 [422], NOTCH3 [423], PCSK6 [424], EMILIN1 [425], CELSR1 [426], IL18 [192], JPH2 [427], CDKN2B [428], GAS6 [429], CMKLR1 [430], THBS1 [431], CBS (cystathionine beta-synthase) [432], CYGB (cytoglobin) [433], EGF (epidermal growth factor) [434], DIO2 [435], TRPM2 [436], THBS2 [174], NOX4 [437], MYLK (myosin light chain kinase) [438], MYH9 [209], COL1A1 [439], GLIS2 [132], MFGE8 [440], TRPV4 [133], PEAR1 [441], CACNA1C [442], LRP5 [443], FLNA (filamin A) [444], GLI1 [445], PIM1 [446], MDK (midkine) [447], TRPC6 [448], PIK3R5 [449], POSTN (periostin) [450], CYP2D6 [451], SDC3 [452], NNMT (nicotinamide N-methyltransferase) [453] and ADAMTS8 [454] functions as a key candidate genes in hypertension. MMP12 [455], BMP2 [456], TAC1 [457], CCK (cholecystokinin) [458], IL33 [459], SORCS1 [460], CALCA (calcitonin related polypeptide alpha) [461], IRF4 [462], CXCL8 [463], MMP3 [464], ABCB1 [465], ADAMTS5 [466], CD24 [467], SPX (spexin hormone) [468], KCNK3 [469], ADCY8 [470], GIPR (gastric inhibitory polypeptide receptor) [471], KCNMA1 [472], CRYM (crystallin mu) [473], PROX1 [474], WNT16 [475], CXCL1 [476], UCN2 [477], PDK4 [478], IL13RA2 [479], SPP1 [480], CXCL5 [481], DNAJC6 [482], DNAJB9 [483], EDNRA (endothelin receptor type A) [484], TXNIP (thioredoxin interacting protein) [485], IL6R [486], ABHD5 [487], HSPA5 [488], DKK1 [489], NAMPT (nicotinamidephosphoribosyltransferase) [490], CREBRF (CREB3 regulatory factor) [491], GPNMB (glycoprotein nmb) [492], GHR (growth hormone receptor) [493], RAG1 [494], SIGLEC15 [495], ANXA1 [496], CD40 [497], HK2 [498], NRG1 [499], CPEB4 [500], PLIN2 [501], CD44 [502], INSIG1 [503], CD47 [504], GYS2 [505], SLC6A15 [506], SAA1 [507], IGFBP2 [508], ELF3 [509], ABI3 [510], PLN (phospholamban) [511], MYF5 [512], MMP9 [513], WNT4 [514], VCAM1 [515], IGF1 [516], CX3CL1 [517], AQP1 [518], NRXN3 [519], ARG1 [520], A2M [521], HTR1B [522], OXTR (oxytocin receptor) [523], SPON2 [524], SULF2 [525], PHOSPHO1 [526], JAK3 [527], LAMA1 [528], IL18 [529], GAS6 [530], CEBPD (CCAAT enhancer binding protein delta) [531], CMKLR1 [532], THBS1 [533], SLC7A8 [534], ARNT2 [535], EGF (epidermal growth factor) [536], DIO2 [537], TRPM2 [538], HSPA2 [539], GPC4 [540], SORT1 [541], NOX4 [542], COL1A1 [340], HSPB7 [543], CPE (carboxypeptidase E) [544], MFGE8 [545], TRPV4 [546], DUSP2 [547], LRP5 [548], C3AR1 [549], RARA (retinoic acid receptor alpha) [550], PHGDH (phosphoglycerate dehydrogenase) [551], PIM1 [552], TRPC6 [553], MACC1 [554], C1QTNF5 [555], ERG (ETS transcription factor ERG) [556], POSTN (periostin) [557], MEST (mesoderm specific transcript) [558], PCSK7 [559], ELOVL2 [560], CYP2D6 [561], STAT4 [562], RCAN2 [563], TM6SF2 [564], AEBP1 [565] and NNMT (nicotinamide N-methyltransferase) [566] were a promising prognostic biomarkers in obesity. MMP12 [567], BMP2 [568], CCK (cholecystokinin) [458], IL33 [569], SORCS1 [570], CXCL8 [571], PENK (proenkephalin) [572], HSD17B3 [573], MMP3 [574], ABCB1 [575], SPX (spexin hormone) [576], NFATC2 [577], ADCY8 [470], ALK (ALK receptor tyrosine kinase) [578], GIPR (gastric inhibitory polypeptide receptor) [579], PODXL (podocalyxin like) [580], KCNMA1 [581], KL (klotho) [582], P2RX7 [583], PROX1 [584], DNER (delta/notch like EGF repeat containing) [585], CXCL1 [586], UCN2 [587], PDK4 [588], CD55 [589], CXCL5 [590], TXNIP (thioredoxin interacting protein) [485], IL6R [591], DKK1 [252], TNFAIP3 [254], CREBRF (CREB3 regulatory factor) [592], PLA2G4A [593], VMP1 [594], LINC-PINT [595], SPRY2 [596], GRB14 [597], GHR (growth hormone receptor) [598], ITGB3 [599], HSPA8 [600], DUSP5 [601], BTG1 [602], ANXA1 [603], CD40 [604], HK2 [498], GLIS3 [605], NRG1 [606], CD44 [607], ARNTL (aryl hydrocarbon receptor nuclear translocator like) [405], EPOR (erythropoietin receptor) [608], VAMP4 [609], INSIG1 [610], CD47 [611], RARRES2 [612], IGFBP2 [613], HSD3B1 [614], ELF3 [615], MMP9 [616], FLT1 [617], VCAM1 [618], IGF1 [619], VTN (vitronectin) [620], CX3CL1 [621], AQP1 [622], ARG1 [623], GNAO1 [624], A2M [625], ACTA2 [636], OAS1 [627], SPON2 [628], SULF2 [525], PHOSPHO1 [629], CDH1 [630], LAMA1 [528], IL18 [631], CDKN2B [632], GAS6 [633], CMKLR1 [634], HES1 [635], CYGB (cytoglobin) [636], ARNT2 [535], EGF (epidermal growth factor) [637], DIO2 [638], TRPM2 [639], TGM2 [640], THBS2 [641], GPC4 [642], SORT1 [643], DACT1 [644], HSPG2 [645], NOX4 [646], MYH9 [647], COL1A1 [648], CPE (carboxypeptidase E) [544], MFGE8 [649], NR2F1 [650], TRPV4 [651], GSN (gelsolin) [652], LAG3 [653], CACNA1C [654], LRP5 [655], RARA (retinoic acid receptor alpha) [656], OAS3 [657], PIM1 [658], TRPC6 [659], POSTN (periostin) [660], CYP2D6 [661], STAT4 [662], TM6SF2 [663] and NNMT (nicotinamide N-methyltransferase) [664] might be involved in the regulation of type 2 diabetes mellitus. MMP12 [665], BMP2 [666], CCK (cholecystokinin) [667], IL33 [668], SORCS1 [669], CXCL8 [670], MMP3 [671], CD24 [672], KCNMA1 [581], KL (klotho) [673], CD274 [674], CXCL1 [675], CLDN4 [676], CD55 [677], CXCL5 [678], TXNIP (thioredoxin interacting protein) [167], CD83 [679], DKK1 [680], TNFAIP3 [681], VMP1 [594], DUSP5 [601], ANXA1 [682], CD40 [683], GLIS3 [607], NRG1 [684], GCLC (glutamate-cysteine ligase catalytic subunit) [685], CD44 [686], CD47 [687], IGFBP2 [688], MMP9 [689], FLT1 [690], CD70 [691], VCAM1 [692], IGF1 [693], AQP1 [694], OAS1 [695], NOTCH3 [696], JAK3 [697], IL18 [698], CYGB (cytoglobin) [636], EGF (epidermal growth factor) [699], TRPM2 [700], BANK1 [701], SORT1 [643], HSPG2 [645], NOX4 [702], MYH9 [703], TRPV4 [704], B4GALNT1 [705], LAG3 [706], CACNA1C [654], LRP5 [707], PIM1 [658], MDK (midkine) [708], CA2 [709], TRPC6 [710], CYP2D6 [711] and STAT4 [712] might be a biological target of type 1 diabetes mellitus. CCK (cholecystokinin) [713], IL33 [714], IRF4 [715], CXCL8 [716], MMP3 [717], DDIT4L [718], KL (klotho) [719], CXCL6 [87], CXCL1 [720], STC1 [721], PDK4 [722], RAB27B [723], CXCL5 [724], GPRC5A [725], TXNIP (thioredoxin interacting protein) [726], RAB3B [727], HSPA5 [728], DKK1 [93], NAMPT (nicotinamidephosphoribosyltransferase) [729], TNFAIP3 [730], GHR (growth hormone receptor) [731], ANXA1 [732], CD40 [733], KITLG (KIT ligand) [734], SMURF2 [735], NRG1 [684], GCLC (glutamate-cysteine ligase catalytic subunit) [685], NPHP1 [736], EPOR (erythropoietin receptor) [737], ADK (adenosine kinase) [738], SAA1 [739], RARRES2 [612], IGFBP2 [613], HSD3B1 [614], ELF3 [615], RUNX3 [740], MMP9 [741], WNT4 [742], FLT1 [743], VCAM1 [744], IGF1 [745], CX3CL1 [746], AQP1 [747], A2M [748], ACTA2 [749], SPON2 [628], RTN1 [750], NOTCH3 [751], IL18 [752], GAS6 [753], THBS1 [754], HES1 [755], CYGB (cytoglobin) [636], EGF (epidermal growth factor) [637], NOX4 [756], MYH9 [757], GLIS2 [758], MFGE8 [759], CD248 [760], RARA (retinoic acid receptor alpha) [761], MDK (midkine) [762], TRPC6 [763], POSTN (periostin) [764], MEST (mesoderm specific transcript) [765], PCDH7 [766], RCAN2 [767], PLEKHH2 [768], AEBP1 [769] and HIC1 [770] promotes the DKD progression. Gene Ontology (GO) and pathways enrichment analysis showed that enriched genes were might be associated with DKD and its complications.

Construction of PPI network and module analysis are easy for investigaters to study the underlying molecular mechanism of DKD and its complications for the reason that the up and down regulated DEGs would be grouped and ordered in the PPI network judging by their interactions. Hub genes related to DKD and its complications were detected owing to Cytoscape. It is speculated that these significant up and down regulated DEGs theoretically lead to the occurrence and progression of DKD and its complications. HSPA8 [174] and MYH9 [209] have been reported to have a essential role in chronic kidney disease. HSPA8 [262], HSP90AA1 [259], VCAM1 [297], MYH9 [339] and PDLIM7 [305] are altered expression in cardiovascular diseases. HSPA8 [174] and VCAM1 [391], MYH9 [209], FLNA (filamin A) [444] and ACTA2 [422] plays an important role in the hypertension. Altered expression of HSPA8 [600], VCAM1 [618] and MYH9 [647] are associated with type 2 diabetes mellitus. HSPA5 [488] and VCAM1 [515] might play an important role in regulating the genetic network related to the occurrence, development of obesity. The abnormal expression of VCAM1 [110] and MYH9 [131] might be related to the progression of renal fibrosis. The abnormal expression of VCAM1 [692] and MYH9 [703] contributes to the progression of type 1 diabetes mellitus. HSPA5 [728], VCAM1 [744] and MYH9 [757] might be related to the pathophysiology of DKD and might be one of the markers for the early diagnosis of DKD. Moreover, SDCBP (syndecan binding protein), HSP90B1, MDFI (MyoD family inhibitor), PML (PML nuclear body scaffold) and DNAJA1 are involved in DKD and its complications, indicating that these novel hub genes might play key roles in the progression of DKD and its complications.

MiRNA-hub gene regulatory network and TF-hub gene regulatory network analyses were used to explore the molecular mechanisms of the hub genes, miRNAs and TFs involved in the occurrence and development of DKD. Research has shown that HSPA8 [174], MYH9 [209], SOX2 [771] and GATA2 [772] plays an important role in the pathogenesis of chronic kidney disease. Studies have revealed that HSPA8 [262], HSP90AA1 [259], MYH9 [339], SORT1 [335], ACTA2 [307], SOX2 [773], SMAD4 [774], GATA2 [775], CCND1 [776], SPI1 [777] and SMARCA4 [778] plays a key role in cardiovascular diseases. The altered expression of HSPA8 [174], MYH9 [209], FLNA (filamin A) [444], SOX2 [779], GATA2 [780], EP300 [781] and SMARCA4 [782] plays a key role in the progression of hypertension. HSPA8 [600], MYH9 [647], CDH1 [630], SORT1 [643], ACTA2 [626], RARA (retinoic acid receptor alpha) [656], hsa-mir-152-3p [783], SOX2 [784] and EP300 [785] are mainly associated with type 2 diabetes mellitus. Studies had shown that HSPA5 [488], SORT1 [541], RARA (retinoic acid receptor alpha) [550], CCND1 [786], EP300 [787] and HOXC9 [788] expression was associated with obesity. HSPA5 [728], MYH9 [757], ACTA2 [749], RARA (retinoic acid receptor alpha) [761], hsa-mir-152-3p [783], SOX2 [789], SMAD4 [790], CCND1 [791] and EP300 [792] were a diagnostic markers of DKD and could be used as therapeutic targets. MYH9 [131], hsa-mir-152-3p [793], SMAD4 [794] and EP300 [792] might be a potential therapeutic target for renal fibrosis treatment. MYH9 [703], SORT1 [643], hsa-mir-301a-3p [795], hsa-mir-125a-5p [796], hsa-mir-375 [797] and SOX2 [798] could be an early detection markers for type 1 diabetes mellitus. We gave a new confirmation for that HSP90B1, HSPH1, LMO2, EED (embryonic ectoderm development), PML (PML nuclear body scaffold), KRT8, hsa-mir-606, hsa-mir-129-1-3p, hsa-mir-5006-3p, hsa-mir-412-3p, hsa-mir-653-5p, hsa-mir-647, CREM (cAMP responsive element modulator) and TCF3 are expected to become a novel biomarkers for DKD and its complications.

## Conclusions

Bioinformatics analysis is a useful tool to explore the molecular mechanism and pathogenesis of DKD and its complications. There were numerous genes that were differentially expressed in the DKD and normal control groups. These hub genes might play key roles in the onset and development of DKD and its complications, and serve as therapeutic targets.

## Acknowledgement

I thank Gualtiero Nagaswaroop Kengunte Nagaraj, Mayo Clinic, Quantitative Health Sciences, 200 1st St SW, Rochester, Minnesota, USA, very much, the author who deposited their NGS dataset GSE217709, into the public GEO database.

## Ethical approval

This article does not contain any studies with human participants or animals performed by any of the authors.

## Informed consent

No informed consent because this study does not contain human or animals participants.

## Availability of data and materials

The datasets supporting the conclusions of this article are available in the GEO (Gene Expression Omnibus) (https://www.ncbi.nlm.nih.gov/geo/) repository. [(GSE217709) https://www.ncbi.nlm.nih.gov/geo/query/acc.cgi]

## Consent for publication

Not applicable.

## Author Contributions

B. V. - Writing original draft, and review and editing

C. V. - Software and investigation

